# From movements to words: action monitoring in the medial frontal cortex along a caudal to rostral prediction error gradient

**DOI:** 10.1101/2024.10.14.618134

**Authors:** Dorokhova Lydia, Shen Shiqing, Peirolo Morgane, Anton Jean-Luc, Nazarian Bruno, Sein Julien, Chanoine Valérie, Belin Pascal, Loh Kep Kee, Runnqvist Elin

## Abstract

Speech error monitoring recruits the medial frontal cortex (MFC) region in the human brain. Error monitoring-related activity in the MFC has been interpreted both in terms of conflict monitoring and feedback-driven control, but as similar regions of the MFC are implicated in various levels of behavioral control ranging from basic motor movement control to high-level cognitive control functions, a more comprehensive account is needed. Moreover, as speech errors and other actions that involve varying control demands engage a widespread yet partially overlapping set of regions of the MFC, such an account should ideally explain the anatomical distribution of error-related functional activations within the MFC. Here we wanted to assess the hypothesis that the MFC has a similar role in the evaluation of action outcomes for motor and mental actions, operating along a rostral-caudal gradient of higher-lower level control demands involving prediction errors from both sensory and epistemic sources. To this end, we conducted an individual-specific annotation of task-fMRI BOLD activation peaks related to speech errors versus correct productions (i.e. that involve the largest cognitive control demands, Study I and II), tongue movement monitoring (i.e. that involve an intermediate level of cognitive and motor control demands) and tongue movement (i.e. that involve only motor control demands, Study II) in the MFC region. Results revealed overlapping clusters across the three contrasts across the MFC, but importantly both the number of peaks and their relative position along the rostral caudal axis were consistent with a hierarchical rostral caudal processing gradient in the MFC. While tongue movement showed more caudal activation in the MFC, speech errors showed more rostral activation, and tongue movement monitoring patterned in between. Furthermore, the combined results of both studies suggested that activation peaks were located more dorsally for participants that had a paracingulate gyrus, replicating a previously documented effect for movement and further supporting a common functional role of the MFC across very distinct actions.

## 1. Introduction

Various actions, ranging from basic movements (Amiez and Petrides (2014); Loh et al. (2020)) to complex cognitive control behaviors (Botvinick et al. (2001)) activate the medial frontal cortex (MFC) in the human brain. Amongst those actions, producing speech errors leads to increased activity in the midcingulate cortex (MCC) (also known as dACC: dorsal anterior cingulate cortex) and in the SMA/preSMA (supplementary motor area/ presupplementary motor area) region (e.g. Runnqvist et al. (2021); Gauvin et al. (2016); Riès et al. (2011); Moller et al. (2006); Todorović et al. (2023)). In current literature, speech error-related MCC activity is often interpreted through the lens of domain-general conflict-based monitoring (e.g., Nozari et al. (2011); Gauvin et al. (2016); Gauvin and Hartsuiker (2020), Riès et al. (2011)). The conflict monitoring theory posits that the MFC, especially the MCC, continuously evaluates conflict levels during cognitive processes. When conflict surpasses a threshold, these structures relay information to other frontal brain regions responsible for executive control, prompting adjustments in processing dynamics. In essence, the occurrence of conflict itself signifies a need for heightened cognitive control, precluding the need of external sensory feedback (e.g., Botvinick et al. (2001)). This theory provides a unified framework for error monitoring during planning and execution, presenting conflict as a continuum with overt errors as its extreme manifestation. A challenge for this theory is that MCC activation, at the heart of conflict monitoring, is robust during error commission but often absent in conflict situations that do not result in overt errors (e.g., Runnqvist et al. (2021); Todorović et al. (2023); Burle et al. (2008)). Involvement of more caudal regions of the MFC, such as the SMA and preSMA has been observed in situations that could be characterized as involving inner conflict (i.e., covert errors) both in the case of speech production and beyond (Alario et al. (2006); Todorović et al. (2023); Bonini et al. (2014)). However, while these situations clearly indicate a role of the caudal MFC regions for performance monitoring, the specific manipulations involved in the studies were measured near articulation or as electromyographic (EMG) responses related to button presses. Therefore, explanations in terms of proprioceptive feedback used for the purpose of performance monitoring cannot be excluded. Similarly, the conflict monitoring account struggles to explain why the same MCC regions activated by speech errors are also involved in simple movements and explicit feedback processing without conflict. Moreover, the exact location of activation related to simple movement and external feedback processing is modulated in a similar way by inter-individual variations in MCC sulcal morphology, suggesting that it is the same region in the MCC, in particular the anterior rostral cingulate motor zone (RCZa) (Amiez and Petrides (2014); Loh et al. (2018); Loh et al. (2020)). Specifically, some individuals have a single cingulate sulcus (CgS) in the MCC, while others have two sulci − a CgS and a paracingulate sulcus (PCgS) which runs dorsally and in parallel to the CgS, forming a paracingulate gyrus between the two sulcus. In individuals with a PCgS, movement and external feedback processing activation align with the PCgS, whereas in those with only a CgS, activation is confined to the CgS. To account for these patterns, an alternative perspective is that of feedback-related control in the MCC. In the case of speech, this account posits a connection between human speech monitoring and vocal feedback monitoring, drawing parallels across human and non-human primates (Procyk et al. (2016); Loh et al. (2018);Loh et al. (2020); see also Runnqvist et al. (2021)). For instance, Loh and colleagues have argued that the pars opercularis of the inferior frontal gyrus (IFG), also known as area 44, governs cognitive control over orofacial and non-speech vocal responses in primates. Meanwhile, the MCC specializes in analyzing vocal non-speech feedback, driving adaptive responses. For the uniquely human control of speech vocal production, the pars triangularis of the IFG (area 45) and pre-SMA are engaged on top of the basic IFG-MCC primate vocal network. Evidence in support of this vocal feedback control network has mainly arisen from studies providing external vocal feedback (Loh et al. (2020)). Recent findings have identified a similar network (BA 44, 45, pre-SMA, MCC) during overt speech errors, but not in high conflict contexts where a correct utterance is produced (Runnqvist et al. (2021); except for manipulations near articulation that could generate sensory feedback Todorović et al. (2023)). This was interpreted as supporting the idea that internal or self-provided feedback, generated through proprioceptive and/or acoustic signals of self-produced speech, drives MCC activation related to speech errors (see also Dorokhova et al. (in press)). However, this perspective can only explain how high conflict in the absence of overt errors triggers activation in regions like the SMA and preSMA for situations involving a motor response and thus generating sensory feedback. Thus, while both the conflict monitoring account and the feedback control account can successfully account for a share of the findings in the literature, neither fully accounts for all actions that elicit activation in overlapping MFC regions. Collectively, the wide range of actions eliciting overlapping MFC activation suggests a shared functional network for a broad set of self-produced actions, from movements to abstract reasoning (Botvinick et al. (2004);Alexander and Brown (2011);Zarr and Brown (2016)). One proposal that could presumably accommodate the full range of actions eliciting MCC activation as well as their precise anatomical distribution is that the MFC acts as a predictor or evaluator of action outcomes, operating along a processing gradient with caudal regions (SMA, preSMA) at one extreme and rostral regions (BA8m, BA9m) at the other (Bonini et al. (2014); Zarr and Brown (2016)). Within the prediction error framework (Zarr and Brown (2016); Alexander and Brown (2011)), the MFC detects discrepancies between observed outcomes and internal rules, with committed errors representing direct mismatches that prompt MFC activation. Additionally, Zarr and Brown (2016) suggests MCC activation during feedback, rather than actual performance, is associated with expectation adjustment and updating. Moreover, Zarr and Brown (2016) delve into the concept of various prediction error signals within the MFC, highlighting their research findings that suggest a hierarchical organization extending from rostral to caudal regions, with distinct subregions specializing in the processing of specific categories of prediction errors. Zarr and Brown (2016) identifies three types of prediction errors: performance errors, high-level prediction errors, and low-level prediction errors. Performance errors occur when an individual makes a mistake due to an inadequate understanding of the current environment, while prediction errors occur when an outcome is different from what was expected. High-level prediction errors occur when an individual’s expectations about the task or environment are violated, while low-level prediction errors occur when an individual’s expectations about sensory input are violated. This proposition has the capacity to encompass all the distinct actions resulting in increased MFC activation, thus including movements, feedback processing, and speech errors. Additionally, this framework’s ability to offer a unified explanation for all the previously mentioned actions indirectly reinforces the concept that basic feedback-control circuitry, initially utilized for movements, vocalizations and other actions involving sensory goals (e.g., Floegel et al. (2023)), has been repurposed to optimize more complex actions, such as language production, involving epistemic goals (Pezzulo et al. (2022)). Thus, while the hierarchical prediction error gradient proposal of MFC function arose from studies on cognitive conflict, it could easily extend to motor actions. In fact, several frameworks of action control or performance monitoring such as predictive coding and internal modeling have highlighted that predictive processes, typically associated with higher order cognitive function, likely evolved gradually from basic sensory-motor loops (e.g., Wolpert and Flanagan (2001); Pezzulo et al. (2022)). This study aims to further develop this continuity hypothesis by providing a detailed, individual-specific characterization of activation peaks within the MFC for actions of varying control demands: speech errors versus correct productions (Study I and II, high-level cognitive control), tongue movement monitoring (Study II, intermediate level cognitive-motor control), and tongue movement (Study II, basic level motor control). First, we will detail preSMA and MCC activations related to speech errors and examine if the effect of sulcal morphology on functional activation location, which was documented for movement control, extends also to speech errors (Study I). Second, we will compare speech error peaks associated with movement and movement monitoring within the same participants (Study II) across a larger region within the MFC (i.e. from SMA to BA9m) to directly assess the presence of a rostral-caudal gradient of control demands.

## 2. Study I

Study I aims to investigate the neural underpinnings of speech error monitoring by focusing on activations in the preSMA and MCC. The primary objective is to determine whether the relationship between cingulate sulcal morphology and functional activations in the MCC, previously documented for movement control, extends to speech error monitoring. By performing a fine-grained, individual-specific characterization of activation peaks within these regions, this study seeks to elucidate the overlapping neural mechanisms involved in speech errors, movement, and feedback processing.

## 3. Methods

The study received appropriate ethical approval (filed under id “EudraCT: 2015-A00845-344” at the regional ethical committee “Comité de Protection des Personnes Sud Méditerranée I”).

### 3.1. Participants

The full set of participants from Runnqvist et al. (2021) were retained for analyses. In that study, twenty-eight (18 females, 10 males) right-handed native speakers of French participated in exchange for monetary compensation. Four participants (4 males) were excluded from the analyses: three because of excessive head movements during the acquisition and one because of a misunderstanding of the task. The average age of the remaining 24 participants was 23,8 (SD 3,2). No participant reported any history of language or neurological disorders.

### 3.2. Task protocol

Participants performed an error-eliciting word pair production task (SLIP, spoonerisms of laboratory induced predisposition, Runnqvist et al. (2021)) in a single session while their bold activity was recorded in an event related protocol inside of an fMRI scanner.

### 3.3. Stimuli and design

Target stimuli consisted of 160 French nouns combined into 80 pairs (the same stimuli as used in Runnqvist et al. (2016)). During the experiment, three priming word pairs preceded each target word pair. The first two shared the initial consonants, and the third pair had further phonological overlap with the error being primed (*sun mall – sand mouth – soap mate – mole sail*). To induce errors, the order of the two initial consonants (/s/ and /m/) is different for the primes and the target. Participants were also presented with 140 filler pairs that had no specific relationship to their corresponding target pairs. One to three filler pairs were presented before each prime and target sequence. Each participant was presented with 460 unique word combinations (80 targets, 240 primes and 140 fillers). Each participant completed six experimental runs in which word pairs were repeated three times in different orders.

### 3.4. Procedure

Participants were instructed to silently read the word pairs as they appeared, naming aloud the last word pair they had seen whenever a question mark was presented, and before the appearance of an exclamation mark. Stimulus presentation and recording of productions to be processed off-line were controlled by a custom-made presentation software compiled using the LabVIEW development environment (National Instruments). Word pairs remained on the screen for 748 ms. Words presented for silent reading were followed by a blank screen for 340 ms. All targets and 40% of the filler items were followed by a question mark for 544 ms, replaced by an exclamation mark presented 544 ms after the presentation of the question mark and remaining for 1020 ms. Before the next trial started there was a blank screen for 544 ms in the case of filler production trials, and jittered between 544 and 1564 in the case of target production trials. The jittered inter stimulus interval was generated according to an exponential function and randomized across runs.

### 3.5. Data

Transcriptions of participants’ spoken productions as well as anatomical images (spatially normalized to the avg152 T1 −weighted brain template defined by the Montreal Neurological Institute (MNI) using the default parameters of nonlinear transformation) and functional images (first-level) corresponding to the contrast of the response accuracy (committed errors vs. correct productions) from Runnqvist et al. (2021) were used in this study. For detailed information regarding data acquisition and data processing, please see Runnqvist et al. (2021).

### 3.6. Analyses

#### Anatomical images

T1-weighted anatomical scans were annotated manually for the presence of a paracingulate sulcus (PCgS) in the left and right hemisphere separately: three labels were assigned: “prominent”, “emerging” and “absent”. The “prominent” PCgS label accounted for the appearance of two deep parallel sulci longer than 40 mm, the “absent” label was assigned for the presence of only one deep cingulate sulcus in the ACC (Figure B.1). The “emerging” label was assigned to group ambiguous cases that did not satisfy the conditions of two previous categories: the upper sulcus was discontinued or not long or laterally deep enough to be classified as such. The examples of several “emerging” cases are given in Figure B.2. 5 participants were observed with no PCgS neither in the left (LH) nor in the right (RH) hemispheres, additionally 8 were found with no right PCgS; the total number of absent cases was therefore 18 hemispheres. 8 participants were observed to have left “prominent” PCgS, and 6 participants were observed to have right “prominent” PCgS, resulting in a total of 14 “prominent” cases. 11 participants were found to have “emerging” PCgS in the left hemisphere and 5 in the right hemispehere, resulting in 16 hemispheres. The sagittal view of the anatomical scans of all subjects can be found in Figure B.4 in Appendix B. For the group analysis of the data, all right hemispheres were flipped along the y-axis to become left hemispheres (using the *flipud* expression of the imCalc function of SPM (SPM Development Team (1991)) on MATLAB R2018b (Inc. (2020)). This was done to increase the number of observable cases and with the assumption that the effect of MCC is bilateral (Runnqvist et al. (2021)). After that, all hemispheres were assigned to the three groups based on their cingulate sulcal morphology: (1) prominent PCgS (14 hemispheres); (2) absent PCgS (18 hemispheres); (3) emerging PCgS (16 hemispheres). As a next step, mean anatomical scans for each group were created.

#### Functional images

##### Individual annotations

We performed individual participant annotations to investigate in detail how the speech error monitoring activations that were observed in the medial frontal cortical region at the group level in (Runnqvist et al. (2021)) were distributed in individual brains. Moreover, we wanted to assess within each participant whether the absence or presence of a PcGS would impact the exact location of functional peaks. Speech error related peaks were examined in both hemispheres of each participant in a region of interest extending from the anterior commissure to the genu of the corpus callosum. Activation peaks were identified based on voxel-level and cluster-level significance: based on the method of Worsley et al. (1996), the statistical threshold for reporting an activation peak voxel as significant (*p* < 0.05) was *t* = 3.57 in a directed search within the medial frontal region (estimated volume = 20000mm3; http://www.bic.mni.mcgill.ca/users/noor/brain_volume.html). At the cluster level, within the same search region, a cluster of voxels with a *t* -value > 2 and a cluster extent > 231mm3 (15 voxels) constituted a significant activation (*p* < 0.05), corrected for multiple comparisons (Friston, Holmes et al; 1995). Within significant clusters, the voxel with the highest *t* -value was identified as the peak activation.

##### Group level analyses

We also performed separate group level analyses on the absent, emerging and prominent PcGS groups with the aim of assessing the location of MCC activation peaks within in group. For this, an identical flipping procedure of the right hemispheres as that applied to the anatomical images was performed on the first-level contrast statistical maps corresponding to contrast of *errors versus correct* trials using the *flipud* expression of the imCalc function of SPM (SPM Development Team (1991)) on MATLAB R2018b (Inc. (2020)) for each participant. As a consequence, all functional activations were located on the left. Subsequently, we implemented a masking procedure, wherein all values corresponding to the right hemisphere were set to zero. This step was undertaken with the objective of exclusively scrutinizing clusters situated within the left hemisphere. We constrained our region of interest for significant peaks to the left hemisphere, extending laterally only up to *x* = −15 in the MNI space. The posterior and inferior boundaries were defined based on coordinates obtained from a reference source BioImage Suite Web (2023), specifically for Brodmann area 24 (MNI coordinates: *y* = −12, *z* = 5). 1-sample t-test analysis of variance was performed on each group (“prominent PCgS”, “absent PCgS” and “emerging PCgS”) separately on the contrast statistical maps corresponding to errors versus correct trials through the Statistical Parametric Mapping software (SPM Development Team (1991)) on MATLAB R2018b (Inc. (2020)).

The resulting level-2 *t* statistic images in each of the three groups were thresholded using the minimum given by a Bonferroni correction and random field theory to account for multiple comparisons (Worsley et al. (1996)). Statistical significance for the group analyses was assessed based on peak thresholds in directed search and the spatial extent of consecutive voxels, just as for the individual analyses. For a single voxel in a directed search, involving all peaks within an estimated gray matter of 300 cm3 covered by the slices, the threshold for significance (*p*< 0.05) was set at *t* = 5.71 for the “Prominent PCgS” group (n=14), at *t* = 5.18 for the “Absent PCgS” group (n=18) and at *t* = 5.39 for the “Emerging PCgS” group (n=16). A predicted cluster of voxels with a volume extent > 70.28 mm^3^ (5 voxels) for the “Prominent PCgS” group (n=14), 68.48 mm^3^ (5 voxels) for the “Absent PCgS” group (n=18), and 69.24 mm^3^ (5 voxels) for the “Emerging PCgS” group (n=16) with a t-value > 3 was significant (*p* < 0.05), corrected for multiple comparisons. All subsequently discussed peak activations were retrieved within the aforementioned delimited region of interest in the SPM12 results tables, all other activations within the significance thresholds but outside the area of interest are reported in Table A.4 in the Annexes.

## 4. Results

### Behavioral results

Out of the 5760 target trials across all participants, 706 resulted in errors (12.3%, MSE 0.4, sd 32.8). The proportion of errors in each morphological group was similar: the Prominent PCgS group produced errors on 13% of the trials, the Absent PCgS group on -12% of the trials and the Emerging PCgS group on -2% of the trials.

### Individual annotations results

Individual subject analyses confirmed the observation at the group level (Runnqvist et al. (2021)) that the speech error monitoring activation in the medial frontal region was made up of three main clusters of activation peaks (Fig. 5, Table 2, Appendix A. Tables A1-A3): one cluster in the presupplementary motor area (preSMA) and two clusters in the midcingulate cortex (MCC1 and MCC2). Figure 5 displays the locations of the individual preSMA, MCC1 and MCC 2 peaks in the left and right hemisphere across all subjects, and in the three cingulate morphology groupings (PCgS Present, PCgS Emerging and PCgS absent) on the standard MNI152 average brain. The average MNI coordinates of the three peaks in the left and right hemisphere, and across the different cingulate morphology groups are reported in Table 2. The individual MNI coordinates of the preSMA, MCC1 and MCC2 peaks are reported in Appendix A. (Tables A1, A2 and A3 respectively).

**Table 1:** Mean coordinates of preSMA, MCC1 and MCC2 peaks across all subjects (All), in subjects with a prominent paracingulate sulcus (Prominent PCgS), in subjects with no paracingulate sulcus (Absent PCgS), and in subjects with emerging paracingulate sulcus (Emerging PCgS). The x, y, z, coordinates are in MNI stereotaxic space. Individual peaks are identified at a significance threshold of p<0.05, corrected for multiple comparisons within the medial frontal cortex.

|  | All |  |  | Prominent PCgS |  |  | Absent PCgS |  |  | Emerging PCgS |  |  |
| --- | --- | --- | --- | --- | --- | --- | --- | --- | --- | --- | --- | --- |
|  | *x* | *y* | *z* | *x* | *y* | *z* | *x* | *y* | *z* | *x* | *y* | *z* |
| **PreSMA** |  |  |  |  |  |  |  |  |  |  |  |  |
| L | -4.9 | 10.4 | 58.8 | -4.5 | 11.0 | 57.5 | -8.0 | 8.5 | 56.0 | -4.2 | 10.5 | 61.0 |
| R | 4.1 | 11.9 | 58.7 | 7.0 | 17.0 | 60.0 | 3.7 | 11.3 | 55.7 | 3.7 | 10.7 | 61.3 |
| **MCC 1** |  |  |  |  |  |  |  |  |  |  |  |  |
| L | -4.5 | 17.2 | 44.2 | -4.9 | 17.0 | 44.9 | -6.5 | 12.5 | 41.5 | -4.3 | 19.3 | 44.6 |
| R | 5.9 | 16.5 | 42.8 | 4.0 | 16.5 | 48.5 | 6.4 | 15.7 | 42.6 | 6.0 | 18.3 | 39.7 |
| **MCC 2** |  |  |  |  |  |  |  |  |  |  |  |  |
| L | -4.8 | 27.1 | 29.9 | -4.7 | 28.0 | 28.0 | -5.0 | 23.0 | 28.0 | -5.3 | 28.3 | 31.4 |
| R | 7.6 | 26.5 | 30.6 | 7.3 | 28.7 | 33.3 | 8.0 | 26.4 | 30.1 | 7.0 | 25.4 | 29.8 |

**Table 2:**
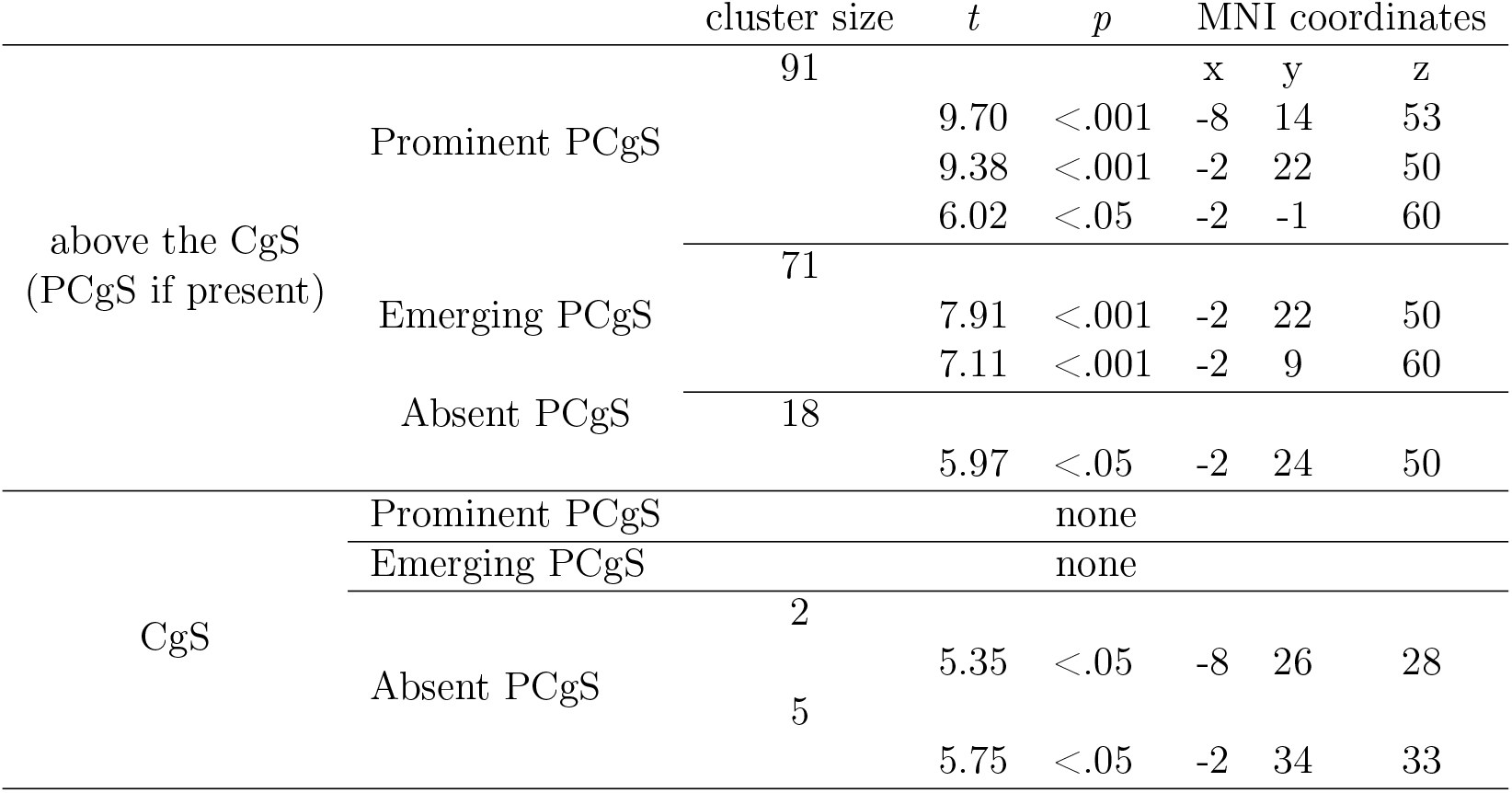
Summary of significant peaks identified in three groups categorized based on their spatial localization, either on the CgS or above it.

The preSMA cluster was consistently located in the medial frontal gyrus just anterior to the VAC line and dorsal to the cingulate gyrus or paracingulate gyrus when it is present (Fig 5). The VAC line is a well-established landmark that separates the supplementary motor area (SMA) posteriorly and the preSMA anteriorly (Ruan et al. (2018)). Across subjects, preSMA peaks were more often observed in the left (14 out of 28 subjects) versus the right (7 out of 28 subjects) hemisphere (Table A1). The average location of the preSMA peak was slightly more dorsal in *PCgS-prominent* (mean MNI *z* -coordinate = 57.9, SD = 4.9) and *PCgS-emerging hemispheres* (mean MNI *z* -coordinate = 61.1, SD = 4.6) than in *PCgS-absent hemispheres* (mean MNI *z* -coordinate = 55.8, SD = 3.2) (Table 2). This suggests that the presence of a paracingulate gyrus (BA32) might be displacing the preSMA region (anterior medial BA6) more dorsally in the medial frontal cortex. As depicted in Brodmann’s classical map (1909), just anterior to the level of the anterior commissure, the anterior part of BA6 (preSMA) lies dorsal to BA32, which constitutes the paracingulate gyrus.

Two distinct clusters of activation peaks were observed across subjects in the midcingulate region extending between the VAC line to the rostral-most limit of the genu of the corpus callosum (Fig 5). This echoes previous work that had similarly revealed the presence of two distinct activations in the same region in behavioral feedback monitoring tasks (e.g. Amiez et al. (2013); Loh et al. (2020)). The first MCC cluster (MCC1) was situated more posterior and dorsal relative to the second MCC cluster (MCC2) and both clusters could be separated by a vertical line drawn from the posterior limit of the genu of the corpus callosum (VPG line; Fig 5). Another crucial observation was that while the MCC2 peaks tend to situate on the cingulate sulcus, regardless of whether a paracingulate sulcus was present, MCC1 peaks appear to situate on the paracingulate sulcus when it is present, and on the cingulate sulcus when a paracingulate was absent (Fig 5). In line with this observation, the average MNI *z* -coordinate of MCC1 peaks were higher in hemispheres with a prominent (mean MNI *z* -coordinate = 45.7, SD = 3.9) and emerging paracingulate sulcus (mean MNI *z* -coordinate = 43.1, SD = 5.1), compared to hemispheres without a paracingulate sulcus (mean MNI *z* -coordinate = 42.3, SD = 4.1)(Table XX). Based on their anatomical locations, MCC1 could be in medial prefrontal area BA8 or cingulate area BA32, and MCC2 could be located more anterior and ventral part of BA32 or BA24 (Brodmann 1909). Lastly, we observed that while MCC1 peaks were more prevalent in the left (17 out of 24) versus the right hemisphere (12 out of 24 subjects), MCC2 peaks are comparably prevalent across the two hemispheres (i.e. 15 versus 17 out of 24 subjects in left and right hemispheres respectively) (Appendix A Tables A2 and A3).

### Group level results

All three group-level analyses yielded significant clusters of activation within the MCC region. The summary of these findings is available in Table 2. The “Absent PCgS” group showed two significant peaks aligned to the CgS that were not present in the other two groups. The remaining activation patterns exhibited a spatial overlap across groups, albeit with quantitative differences. The “Prominent PCgS” group displayed three significant peaks of activation within a substantial cluster measuring 1419.6 mm^3^, situated over the PCgS and extending along the anterior-posterior axis from VAC to VPG. The lateral extent of these peaks ranged from *x* = −2 to *x* = −8 (MNI coordinates).

Conversely, the “Emerging PCgS” group also demonstrated two significant peaks of activation within the same region, specifically above the CgS. The anterior-posterior extent of this cluster mirrored that of the “Prominent PCgS” group, spanning from VAC to VPG. However, the lateral extent did not reach *x* = −8; both peaks were significant at *x* = −2 (MNI coordinates). Furthermore, the cluster size for the “Emerging PCgS” group was smaller, measuring 1107.6 mm^3^.

Lastly, the “Absent PCgS” group exhibited a significant peak within a notably smaller cluster measuring 280.8 mm^3^, also located above the CgS. However, both the anterior-posterior and lateral extents of this cluster were more limited than two other groups.

## 5. Interim discussion

The results of the individual analyses of our Study I indicate that speech errors elicit activation in three distinct clusters in the MFC: one in the pre-supplementary motor area (preSMA), and two peaks in the midcingulate cortex that differed in position both along the rostral-caudal axis and along the dorsal-ventral axis. These results are very similar to those reported in the whole brain analysis in Runnqvist et al., but add information about laterality and reliability across individual participants. Moreover, the first of these two MCC peaks varied in position depending on participants sulcal morphology. Participants with a prominent PCgS showed activations primarily on the PCgS itself, while those with an emergent PCgS showed activations above the cingulate sulcus (CgS). Conversely, participants without a PCgS showed activations on the CgS itself, with lesser activations in the PCgS region. The preSMA cluster showed a similar pattern of activation in function of the presence or absence of a PcGS. This pattern was complemented by the group analyses, showing that a cluster extending from the SMA to 8m was increasingly more dorsal and larger with increased presence of a PcGS; and that a cluster located on the CGS in 8m extending to 9m was only present for the group lacking a PcGS. Interestingly, the three distinct clusters of activation in the MFC that were observed for speech errors in the present study bear a high similarity with those reported in a previous study (**?**). (**?** used a task involving sorting objects based on regularly modified rules to induce prediction errors. The results showed that the medial prefrontal cortex (mPFC) responded differently to the three types of prediction errors, with precise spatial organization and functional connections with the lateral prefrontal cortex. Posterior regions of the mPFC were associated with less abstract prediction errors, while anterior regions were associated with more abstract prediction errors. These findings support the idea of a rostro-caudal gradient of abstraction in the mPFC. Returning to our findings, as language is a hybrid action, entailing both motor and cognitive control, one possibility is thus that the three clusters reflect the engagement of control that is more or less motor related along a hierarchical gradient. This hypothesis will be tested in our second study.

## 6. Study II

The aim of this study was to provide a fine grained characterization of speech error, tongue movement monitoring and tongue movement related MFC activations. For each participant, peaks were categorized into 12 categories along the rostral-caudal and dorsal-ventral axes. We extended the search space area to include the SMA on the caudal extreme and medial prefrontal area 9m on the rostral extreme of the medial frontal cortex. This decision was driven by the fact that in the results of study one, the clusters at the caudal and rostral extremes were at the border in between SMA and preSMA, and between medial prefrontal areas 8m and 9m respectively. This suggests that there were in fact more than 3 relevant categories along the rostral caudal axis. Similarly, along the dorsal ventral axis, the 3 clusters that we observed in study 1 all differed along this dimension, hinting at the interest of exploring the dorsal ventral dimension along the different rostral caudal categories.

### 6.1. Methods

The study received ethical approval from the regional ethical committee, Comité de Protection des Personnes Sud Méditerranée I, under identification number 2017-A03614-49.

### 6.2. Participants

Twenty right-handed young adults, including 12 women, participated in the study, with ages ranging from 20 to 33 years (mean age = 25.5 years). Participants were compensated for their participation. None of the participants reported a history of language or neurological disorders.

### 6.3. Task protocol

Participants took part in four experimental sessions separated by a week. In all four sessions they performed an error eliciting language production task similar to that of Study I, and in the first session only they also performed a task of movement and movement monitoring (see description below). For the language production task, only the aspects that differed from Study I will be described.

### 6.4. Stimuli and design

#### Language production task

Target stimuli consisted of 112 French nouns combined into 56 pairs. During the experiment, 3 phonological prime word pairs preceded each target word pair. In half of the target pairs, the primed error outcome of the first word was additionally primed with a synonym placed after the phonological primes (e.g., “trace pose” for “barque manque,” where “trace” is a synonym for “marque”). 53% of the filler pairs were presented for silent reading. Each participant was presented with 410 unique word combinations (56 targets, 224 primes and 130 fillers), and this was repeated twice in each experimental session, with a different order, partitioned into four runs, each lasting for 5 minutes. Each participant engaged in 4 experiment sessions (16 runs), with a span of one week between each session. *Movement task*. Target stimuli consisted of written words (hand, tongue, fixate) in white or red font indicating the action to be performed, and a fixation cross, indicating that the action should be performed.

### 6.5. Procedure

#### Language production task

Word pairs remained on the screen for 693 ms. Words presented for silent reading were followed by a blank screen for 231 ms. All targets and 40% of the filler items were followed by a question mark for 539 ms, replaced by an exclamation mark remaining for 1001 ms. Before the next trial started there was a blank screen jittered between 462 and 1232 for both filler and target production trials. The jittered inter stimulus interval was generated according to an exponential function and randomized across runs.

#### Movement task

Participants were instructed to look at a centrally presented fixation cross and either produce recurrent tongue movements, recurrent hand movements or no movement at all (baseline, e.g., Amiez and Petrides (2014)). The condition was indicated by a printed word (tongue, hand or fixate) presented for 1001 ms prior to the appearance of the fixation cross. The fixation cross remained for 15015 ms on the screen. For hand movements, participants made up and down movements with their wrist and hand (fingers straightened) using their working (right) hand. For tongue movements, participants made vertical circular movements with their tongue, keeping their lips closed but loosening their jaws to create more space in the oral cavity. For both types of movements, participants were instructed to make one movement cycle per second: up and down or a circle. For both body parts there were two movement conditions: the movement alone (described above) and the monitored movement). For the monitored movement condition, participants were given additional instructions not to touch the bed of the fMRI scanner for hand movements, and not to touch the walls of the oral cavity or teeth for tongue movements. To differentiate between the movement alone condition and the monitored movement condition, the former was displayed in white and the latter in red font. One run contained 2 repetitions of each of the 5 conditions (10 trials per run: 2 hand movement, 2 hand monitored movement, 2 tongue movement, 2 tongue monitored movement, 2 fixate). Participants completed 4 runs of 10 trials (120 sec per condition overall).

#### 6.5.1. Data Acquisition

A 3-Tesla Siemens Prisma Scanner (Siemens, Erlangen, Germany) equipped with a 64-channel head coil at the Marseille MRI Center (Centre IRM-INT @CERIMED, UMR7289 CNRS & AMU) was utilized to acquire data. Functional BOLD images were collected using an EPI sequence with 72 slices per volume, multi-band = 4, repetition time = 1.386 s, spatial resolution = 2 × 2 × 2 mm3, echo time = 33.4 ms, and flip angle = 56°, covering the entire brain during task performance. Whole-brain anatomical MRI data were obtained using a high-resolution structural T1-weighted image (MPRAGE sequence) with a repetition time of 2.3 s, spatial resolution of 0.8 × 0.8 × 0.8 mm3, echo time of 3.1 ms, inversion time of 0.9 s, and flip angle of 9° in the sagittal plane. Prior to functional imaging, whole-brain Fieldmap images were acquired using a spin-echo EPI sequence with the same spatial parameters as the BOLD images, acquired twice with opposite phase encode directions along the anterior-posterior axis. These images had the following parameters: TR/TE = 7220/59 ms, voxel size = 2 × 2 × 2 mm3, slices = 72, and flip angle = 90/180°.

#### 6.5.2. Behavioral data processing

##### Language production task

A person naïve to the purpose of the experiment transcribed all spoken productions. If participants produced the pair correctly despite the priming, the trial was considered as “correct”; any production that did not match the written word pair was considered as “error”.

#### 6.5.3. Neuroimaging data processing

The data underwent preprocessing using fMRIPrep 20.2.6 (Esteban et al. (2018b) ; Esteban et al. (2018a), a tool built on Nipype 1.7.0 (Gorgolewski et al. (2011) ; Gorgolewski et al. (2018).

#### 6.5.4. Anatomical data preprocessing

The T1-weighted (T1w) image underwent intensity non-uniformity (INU) correction using N4BiasFieldCorrection (Tustison et al. (2010), distributed with ANTs 2.3.3 Avants et al. (2008), and served as the T1w-reference throughout the workflow. Subsequently, the T1w-reference underwent skull-stripping using a Nipype implementation of the antsBrainExtraction.sh workflow (from ANTs), with OASIS30ANTs as the target template. Brain tissue segmentation into cerebrospinal fluid (CSF), white matter (WM), and gray matter (GM) was performed on the brain-extracted T1w image using fast (FSL 5.0.9, Zhang et al. (2001)). Brain surfaces were reconstructed using recon-all (FreeSurfer 6.0.1, Dale et al. (1999)), and the previously estimated brain mask was refined using a custom variation of the method to reconcile ANTs-derived and FreeSurfer-derived segmentations of cortical gray matter from Mindboggle (Klein et al. (2017)). Volume-based spatial normalization to the MNI152NL in 2009 cAsym standard space was conducted through nonlinear registration using antsRegistration (ANTs 2.3.3), utilizing brain-extracted versions of both the T1w reference and template. The ICBM 152 Nonlinear Asymmetrical template version 2009c (Fonov et al. (2009), TemplateFlow ID: MNI152NL in2009 cAsym) was selected as the template for spatial normalization.

#### 6.5.5. Functional data preprocessing

The preprocessing steps for each of the BOLD runs in each task (4 for the movement task, 16 for the language task) per subject included several procedures. Initially, a reference volume and its skull-stripped version were created by aligning and averaging single-band references (SBRefs). A B0-nonuniformity map, or fieldmap, was then estimated using two echo-planar imaging (EPI) references with opposing phase-encoding directions, utilizing 3dQwarp from AFNI 20160207 (Cox and Hyde (1997)). Subsequently, a corrected EPI reference was generated based on the estimated susceptibility distortion, facilitating more accurate co-registration with the anatomical reference. The co-registration process involved aligning the BOLD reference to the T1-weighted (T1w) reference using bbregister from FreeSurfer, which employs boundary-based registration (Greve and Fischl (2009)). Head-motion parameters with respect to the BOLD reference were estimated using mcflirt from FSL 5.0.9 (Jenkinson et al. (2002)), before applying any spatiotemporal filtering. The BOLD time-series, including slice-timing correction if applied, were resampled into their original, native space by applying a single, composite transform to correct for head-motion and susceptibility distortions. Subsequently, the BOLD time-series were resampled into standard space, resulting in preprocessed BOLD runs in MNI152NL in 2009 cAsym space. Additionally, two global signals were extracted within the cerebrospinal fluid (CSF) and white matter (WM), and physiological regressors were extracted to facilitate component-based noise correction (CompCor, Behzadi et al. (2007)).

### 6.6. Analyses

#### Anatomical images

T1-weighted anatomical scans were annotated manually for the presence of a paracingulate sulcus (PCgS) in the left and right hemisphere separately following the same criteria as in Study I. 7 participants were observed to not have a right PCgS and 2 with no left PCgS; the total number of absent cases was therefore 9 hemispheres. 13 participants were observed to have left “prominent” PCgS, and 5 participants were observed to have right “prominent” PCgS, resulting in a total of 18 “prominent” cases. 5 participants were found to have “emerging” PCgS in the left hemisphere and 8 in the right hemispehere, resulting in 13 hemispheres.

#### Functional images

The preprocessed BOLD data were then analyzed using the Statistical Parametric Mapping software (SPM12) on MATLAB R2020a. The BOLD data were smoothed with an isotropic three-dimensional Gaussian kernel (FWHM = 5 mm), and a general linear model (GLM) was designed for each run and subject. In the case of the language task the GLM contained 8 conditions (semantically related distance 1; semantically related distance 3; semantically unrelated distance 1, semantically unrelated distance 3; error; silence; prime and filler). In the case of the movement task the GLM contained 6 conditions (fixation, hand movement, monitored hand movement, tongue movement, monitored tongue movement and instruction). The regressors were convolved with the canonical hemodynamic response, and 48 nuisance regressors were included in the model, comprising 24 head movement-derived regressors, 24 aCompCor regressors (12 CSF and 12 WM), and two global signals within CSF and WM masks. For the present purposes, we used the first level images corresponding to the contrasts of “errors minus correct productions” from the language production task (henceforth referred to as speech error), and to the contrasts of “tongue movement minus fixation” (henceforth referred to as movement) and “tongue movement monitoring minus fixation” (henceforth referred to as monitoring) from the movement task.

We performed individual subject annotations to investigate in detail how the activation peaks corresponding to the three different task contrasts were distributed in individual brains in the medial frontal cortex. Peaks were examined for each contrast and in each subject in the medial frontal cortical region in a search area spanning from the rostral limit of the pons (MNI y-coordinate = -12) to the rostral end of the brain, with a ventral boundary at the limit of medial prefrontal area 9 (MNI z-coordinate = -3) (see Figure 3). Activation peaks were identified based on voxel-level and cluster-level significance: based on the method of Worsley et al. (1996), the statistical threshold for reporting an activation peak voxel as significant (*p* < 0.05) was *t* = 3.57 in a directed search within the medial frontal region (estimated volume = 20000mm3; http://www.bic.mni.mcgill.ca/users/noor/brain_volume.html). At the cluster level, within the same search region, a cluster of voxels with a *t* -value > 2 and a cluster extent > 231mm3 (15 voxels) constituted a significant activation (*p* < 0.05), corrected for multiple comparisons (Worsley et al. (1996)). Within significant clusters, the voxel with the highest *t* -value was identified as the peak activation. Within the region of interest, peaks were classified into 12 distinct regions depending on their position along the rostral-caudal axis (SMA, preSMA/SMA, preSMA, medial BA 8 (“8”), and medial BA 9 (“9”), and along the dorsal-ventral axis (ranging from 1 (most dorsal) to 3 (most ventral; see Figure 3). If there were more than one significant peak in a particular MFC subdivision, the peak with the highest t-value will be selected.

**Figure 1:**
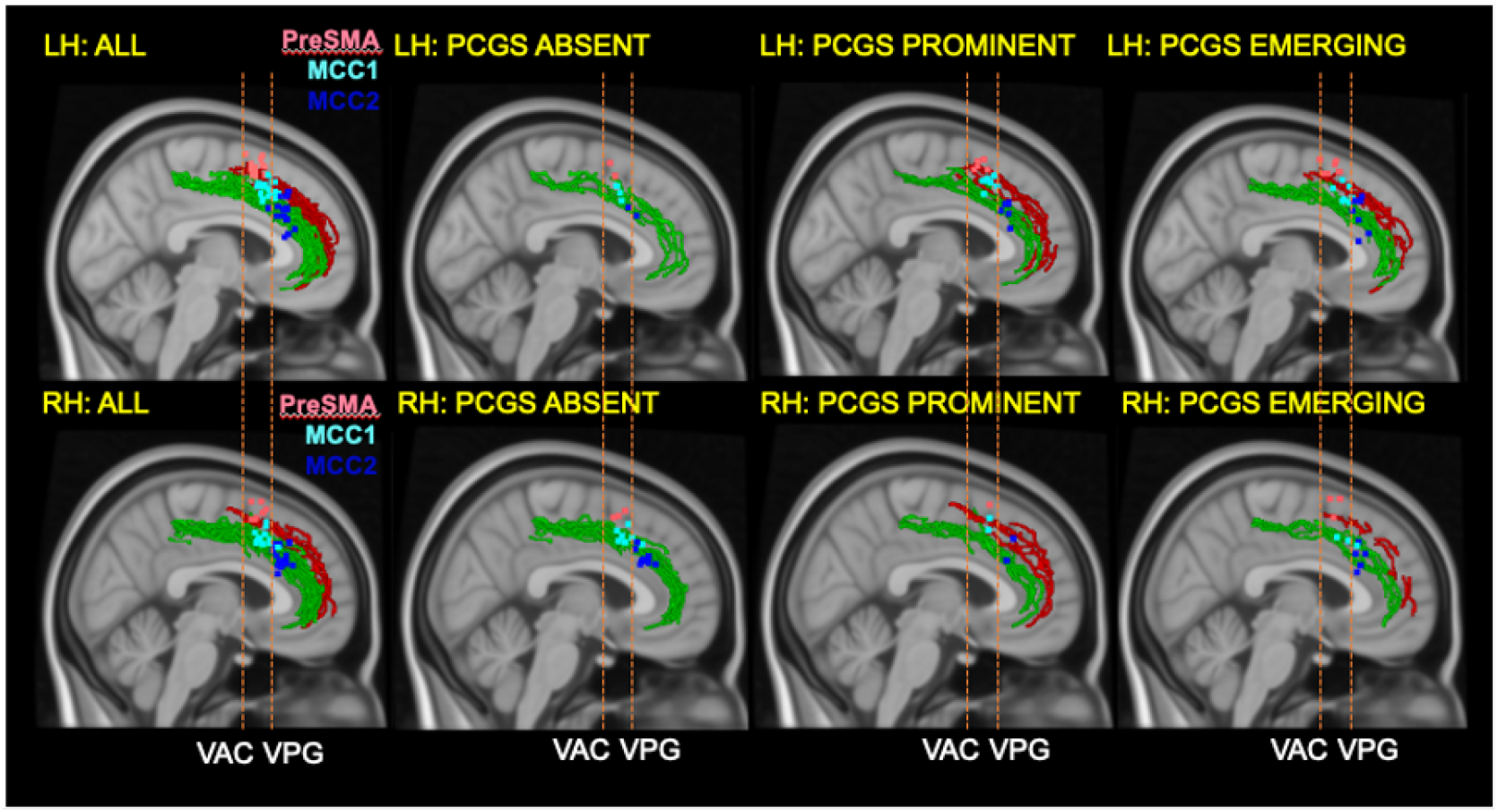
Individual subject analyses of preSMA, MCC1 and MCC2 peaks in relation to cingulate morphology. Individual preSMA (pink), MCC1 (light blue) and MCC2 (dark blue) peaks are depicted on the standard MNI152 brain in the left (Top row) and right hemispheres (Bottom row) across all subjects (All), and for cases without a paracingulate sulcus (PCGS Absent), with a prominent paracingulate sulcus (PCGS Prominent), and with an emerging paracingulate sulcus (PCGS Emerging). Individual peaks are identified at a significance threshold of p<0.05, corrected for multiple comparisons within the medial frontal cortex. VAC=Vertical line from the anterior commissure. VPG=Vertical line from the posterior limit of the genu of the corpus callosum.

**Figure 2:**
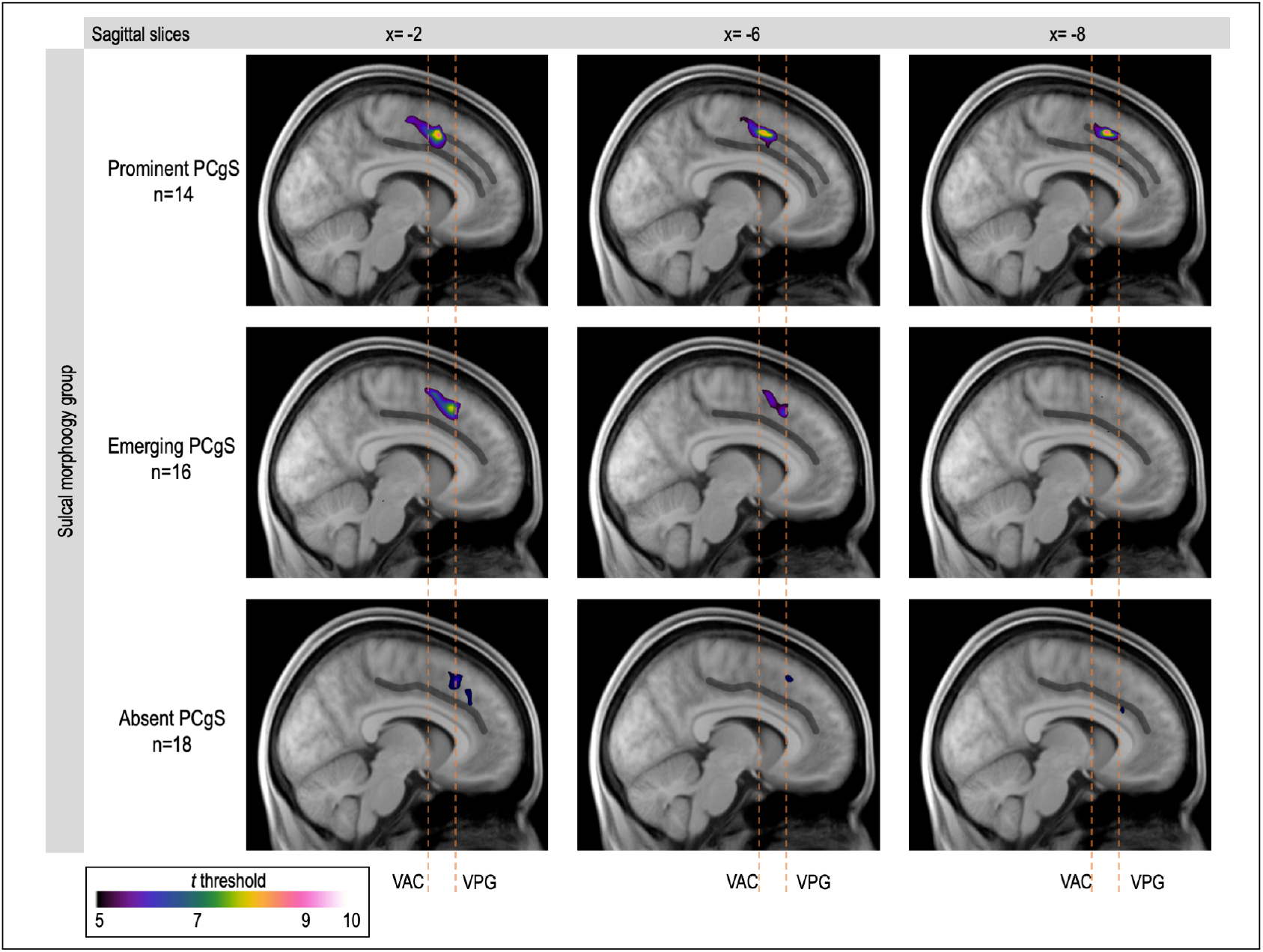
Results for three groups of all right and left hemispheres mixed presented on mean anatomical scans of each group, the CgS and PCgS are drawn in gray lines. VAC refers to the vertical line drawn through anterior commissure, VPG refers to vertical line drawn through the posterior limit of the genu of the corpus callosum

**Figure 3:**
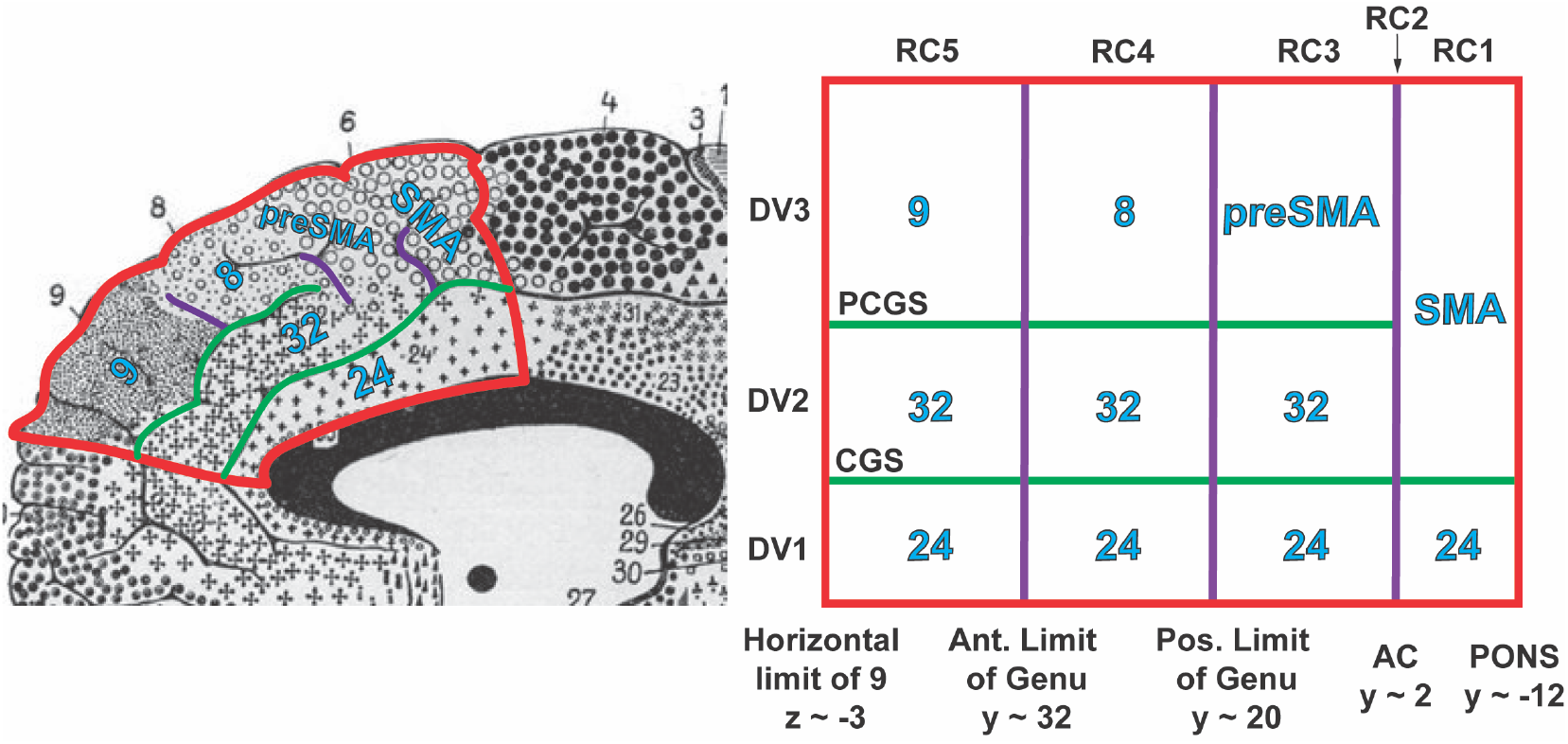
Schematic of the anatomical subdivisions of the medial frontal cortex in relation to anatomical landmarks. Numbers correspond to Brodmann areas. RC1-RC5: caudal to rostral positions within the medial frontal cortex; DV1-RC3: ventral to dorsal positions within the medial frontal cortex; AC=Anterior Commissure; CGS = cingulate sulcus; PCGS = paracingulate sulcus. Note that RC2 refers to the position along the vertical line from the AC.

In first instance, we provide descriptive statistics of the findings of our individual annotations as is customary in the previous literature. Where supported by the amount of observations, the data were subsequently analysed using the lme4 package (Bates et al. (2015)) in R version 3.6.3 (R Core Team 2020). The presence of peaks was analysed using generalized linear mixed models (GLMM) with a binomial link function (e.g., Jaeger (2008)), estimating the conditional probability of a response given the random effects and covariate values. The three dimensions of MNI coordinates (x, y and z) were analysed with linear mixed models (LMM), estimating the influence of fixed and random covariates on the response (e.g., Baayen et al. (2008)). In all models, Participants were included as random effect. The fixed factors (varying depending on the model, see below) were “contrast” with the levels “movement”, “monitoring” and “speech” linearly contrast coded; “rostral.caudal” with the levels “RC1”, “RC2”, “RC3, “RC4” and “RC5” linearly contrast coded; “dorsal.ventral” with the levels “DV1”, “DV2” and “DV3” linearly contrast coded; “hemisphere” with the levels “left and “right”, and “PCgS” with the levels “PCgS-A” (absent), “PCgS-E” (emerging), and “PCgS-P” (prominent) linearly contrast coded. The dependent variables were either the peaks (present or absent) or the three dimensions of MNI coordinates (x, y and z).

The probability of the presence or absence of peaks in function of the fixed variable “rostral.caudal” as a function of the fixed variable “contrast” was assessed using binomial logistic regression (model a). This model also included the fixed variables : 1) Hemisphere (left or right) and 2) Task (movement, monitoring or speech error) and all interactions between the three fixed variables. In a complementary approach, we also fitted linear models of the y coordinates of the peaks to examine the impact of Task and Hemisphere on the rostral-caudal position of peaks from a more continuous perspective (model b). A rostral caudal gradient of task control demands should result in an interaction between Rostral-Caudal and Task with higher levels of task control demands leading to more rostral MFC activations; and in a positive main effect of Task on y coordinates with higher levels of task control demands leading to increased y-values (i.e. more rostral activations). Finally, we also modeled the linear impact of the variable “PcGS” with the levels “absent”, “emerging” or “prominent” on the presence or absence of peaks in each of the 12 MFC subregions and on the z coordinates. Statistical support for the hypothesis that the presence of a PcGs results in more dorsal activation in the MFC would be obtained through a main effect of the PcGS variable on the z coordinates with the increasing presence of a PcGS predicting higher z-values (more dorsal activations).

## 7. Results

### Behavioral results language production task

Out of the 8960 target trials across all participants, 1288 resulted in errors (14.4%, MSE 1,2%, sd 7,5%). The proportion of errors in each morphological group was similar: the Prominent PCgS group produced errors on 14.2% of the trials, the Absent PCgS group on 16.5% of the trials and the Emerging PCgS group on 11.5% of the trials.

### A caudal-to-rostral shift in MFC activations going from basic movement control to movement monitoring to speech error monitoring

We performed a binomial logistic regression (model a) predicting peak probability with the caudal-to-rostral position within the MFC coded as a linear contrast (i.e. RC1 to RC5; see Table 3), hemisphere, and task condition (i.e. movement, movement monitoring and speech error monitoring). The regression was performed using the glmer program from the lme4 package in R via the following command:

*glmer(peak rostral.caudal*hemisphere*contrast+(1|subject), data=data.df, family=binomial(link=‘logit’))*.

**Table 3:** Model a: glmer(peak rostral.caudal*dorsal.ventral*contrast+(1|subject), data=data, family=binomial)

|  | Estimate | Std.Error | z value | Pr(>\|z\|) |
| --- | --- | --- | --- | --- |
| (Intercept) | -0.12872 | 0.12668 | -1.016 | 0.309582 |
| rostral.caudal.L | -0.49142 | 0.18028 | -2.726 | 0.006415 ** |
| hemisphereR | -0.26007 | 0.11372 | -2.287 | 0.022198 * |
| contrast.L | 0.11610 | 0.13855 | 0.838 | 0.402037 |
| rostral.caudal.L:hemisphereR | 0.18801 | 0.25747 | 0.730 | 0.465255 |
| rostral.caudal.L:contrast.L | 1.14355 | 0.31482 | 3.632 | 0.000281 *** |
| hemisphereR:contrast.L | -0.07945 | 0.19809 | -0.401 | 0.688368 |
| rostral.caudal.L:hemisphereR:contrast.L | 0.39237 | 0.45077 | 0.870 | 0.384054 |

There was a significant interaction between task condition and caudal-to-rostral position (see Table 3): As evident from Figure 5, peak probabilities are higher in more caudal MFC locations in the movement condition, but are higher in more rostral MFC locations in the speech monitoring condition. Interestingly, in the monitoring condition, the caudal-to-rostral gradient in peak probabilities was an intermediate between the two other conditions. This trend was also observed in the heatmap plot (Figure 4) with higher percentages of peaks detected in more rostral MFC positions going from movement to monitoring to speech monitoring. In line with our hypothesis, this result indicated that as the complexity of the monitoring increased (i.e. from producing basic tongue movements, to the monitoring of tongue movements, and to the monitoring of speech errors), there was an increased engagement of more rostral parts of the MFC and a decreased engagement of more caudal parts of the MFC.

**Figure 4:**
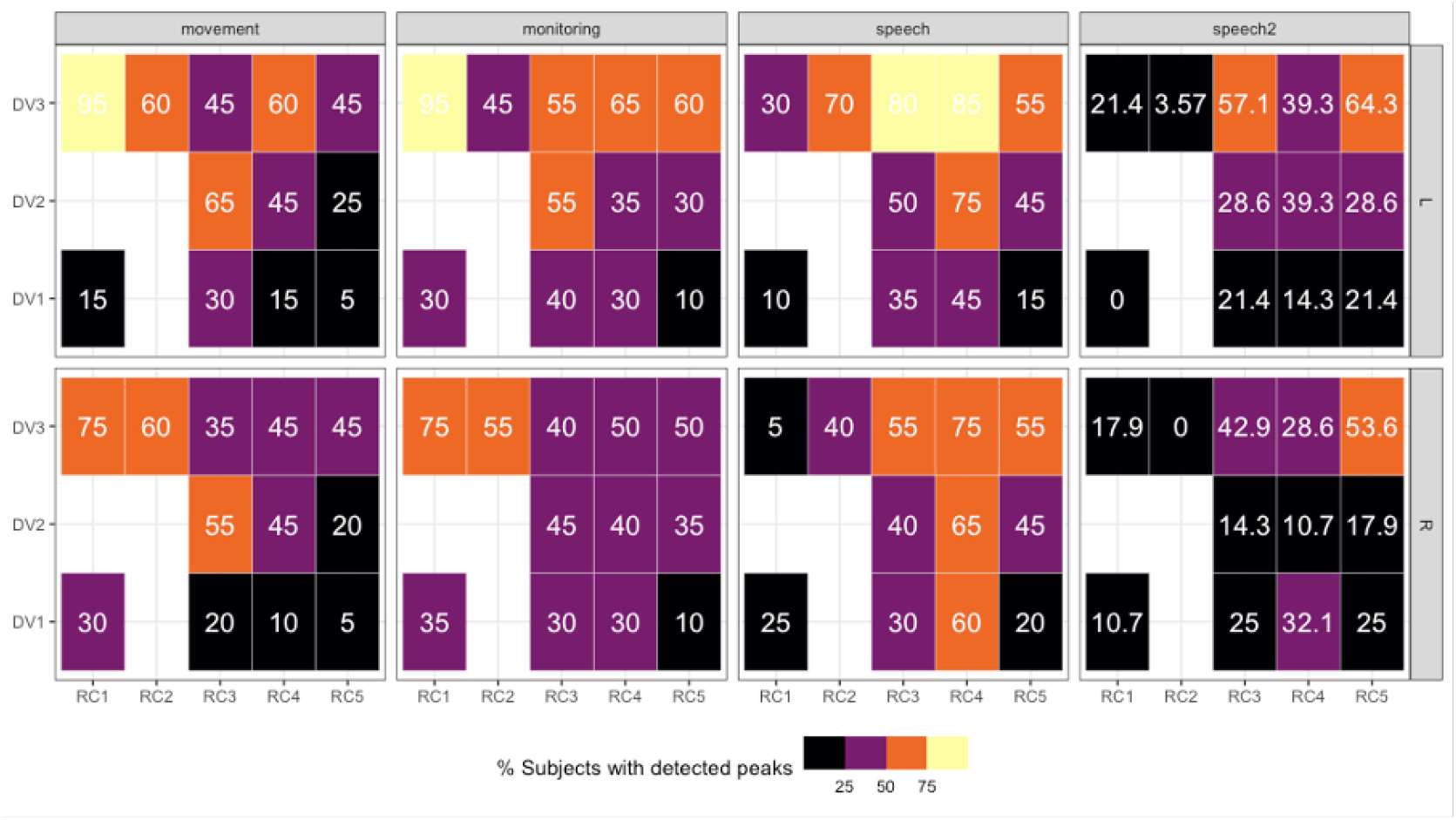
Heatmap showing the proportion of subjects showing a significant peak in each medial frontal cortical subregion for each task condition: movement, monitoring, speech error monitoring (Study2) and speech error monitoring (Study1). RC1-RC5 indicate the caudal-to-rostral position of each subregion. DV1-DV3 indicate the ventral-to-dorsal position of each subregion.

**Figure 5:**
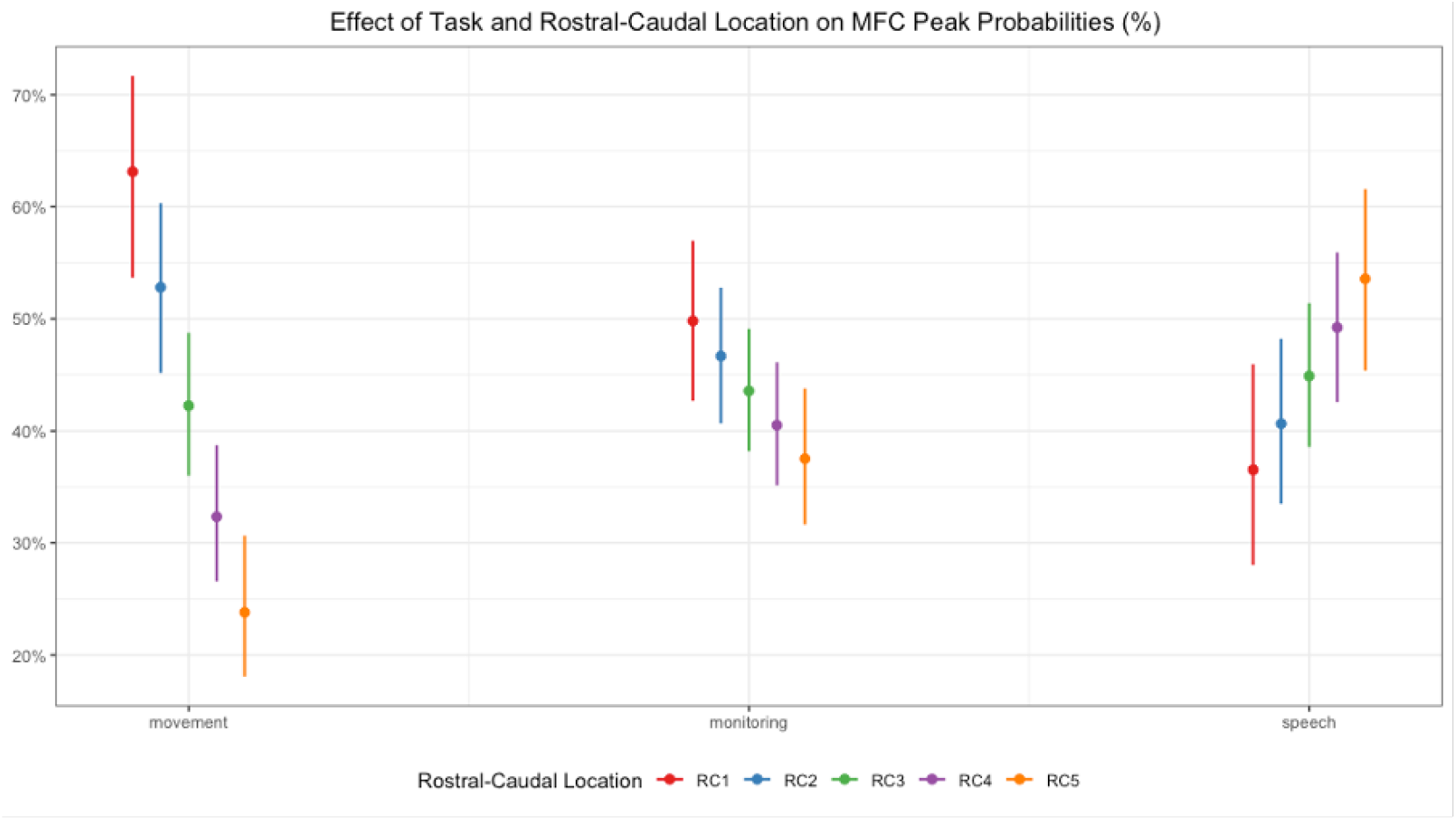
Predicted MFC peak probabilities plotted by rostral-caudal location and task condition. Predicted peak probabilities were obtained via the fitting of model a : glmer(peak rostral.caudal*hemisphere*contrast+(1|subject), data=data.df, family=binomial(link=‘logit’)), followed by the *ggmmeans* function.

From model a, there also was a significant fixed main effect of *hemisphere* on peak probability (see Table 3): the chance of detecting a peak in the left hemisphere was significantly higher than in the right hemisphere across hemispheres and task conditions (Figure 6). This finding suggested that the production and monitoring of tongue movements and the monitoring of speech productions are more localised in the left hemisphere.

**Figure 6:**
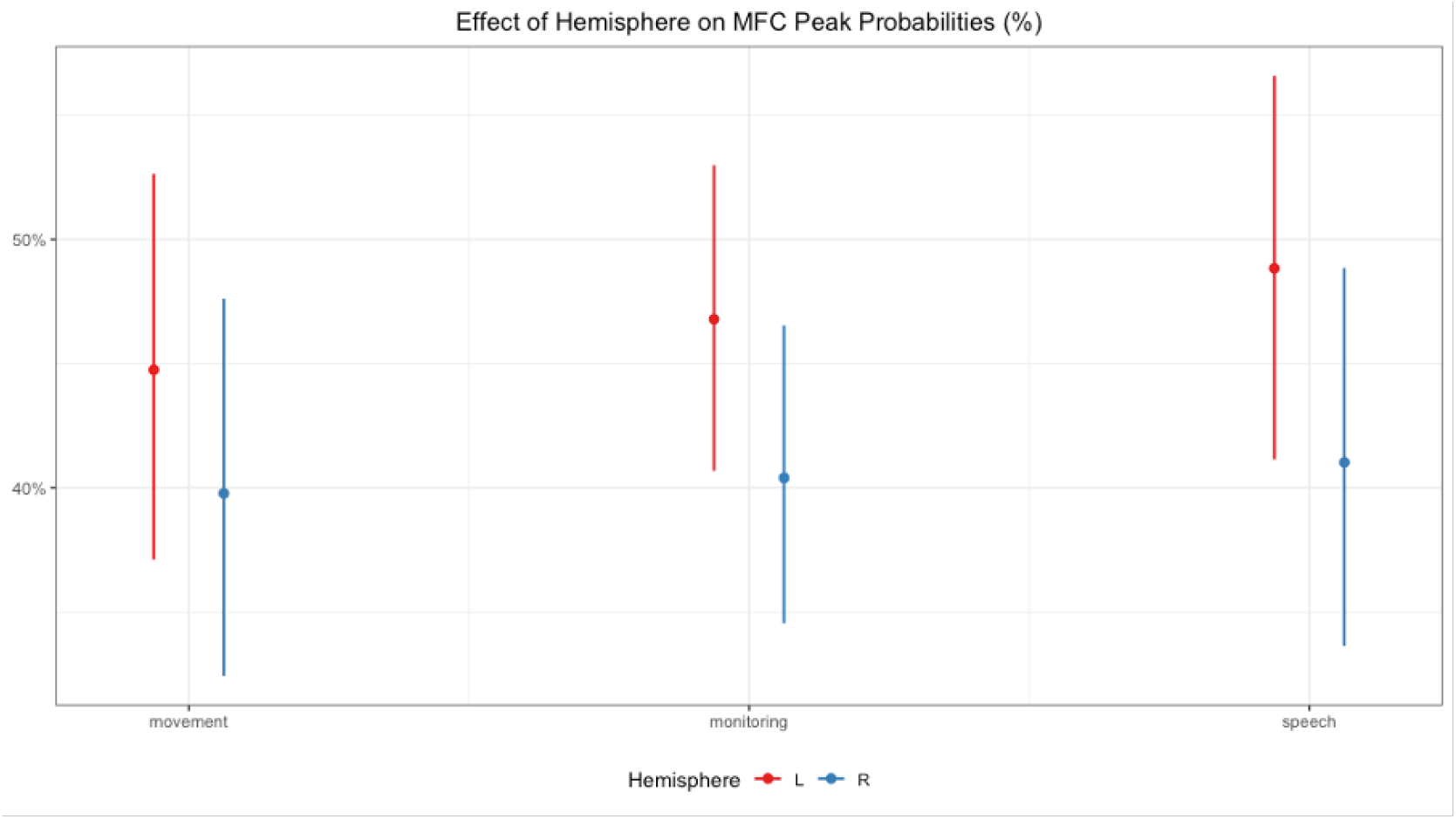
Predicted MFC peak probabilities plotted by hemisphere and task condition. Predicted peak probabilities were obtained via the fitting of model a : glmer(peak rostral.caudal*hemisphere*contrast+(1|subject), data=data.df, family=binomial(link=‘logit’)), followed by the *ggmmeans* function.

In a set of complementary analyses, we performed a general linear model analysis (Table 4) predicting the MNI y-values of the MFC activation peaks with task condition and hemisphere: *lmer(y hemisphere * contrast + (1*|*subject), data=data.df)*. This model yielded a significant fixed main effect of task contrast on the y-values of the MFC activation peaks: Going from basic movement control to movement monitoring to speech error monitoring, the average y-values of the activation peaks in the MFC increased (i.e. shifted more rostrally; Figure 7).

**Table 4:**
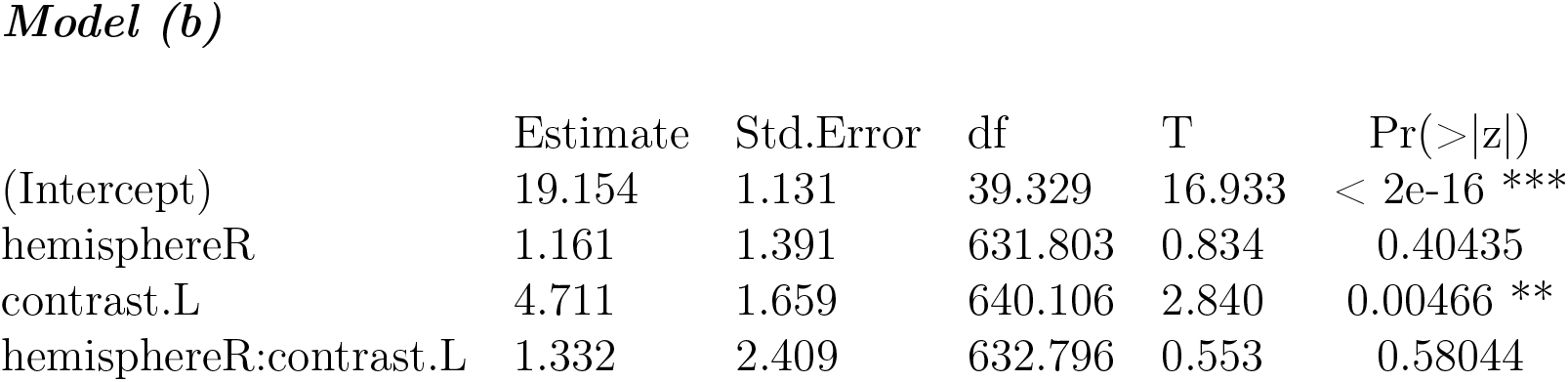
Model b: lmer(y hemisphere * contrast + (1|subject), data=data.df)

|  | Estimate | Std.Error | df | T | Pr(>\|z\|) |
| --- | --- | --- | --- | --- | --- |
| (Intercept) | 19.154 | 1.131 | 39.329 | 16.933 | < 2e-16 *** |
| hemisphereR | 1.161 | 1.391 | 631.803 | 0.834 | 0.40435 |
| contrast.L | 4.711 | 1.659 | 640.106 | 2.840 | 0.00466 ** |
| hemisphereR:contrast.L | 1.332 | 2.409 | 632.796 | 0.553 | 0.58044 |

**Figure 7:**
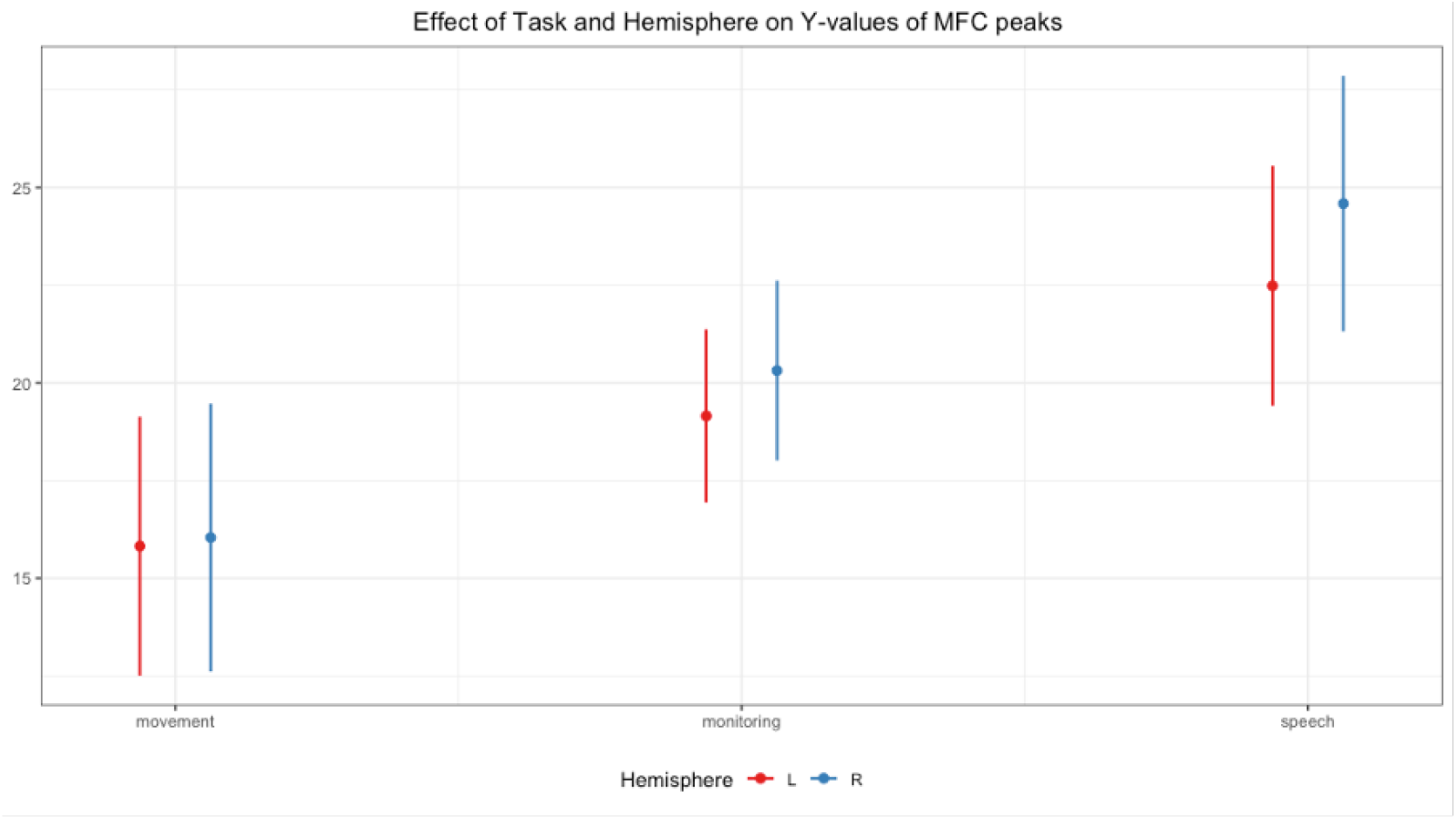
Predicted y-values of MFC activation peaks plotted by hemisphere and task condition. Predicted peak probabilities were obtained via the fitting of model b : lmer(y hemisphere * contrast + (1|subject), data=data.df), followed by the *ggmmeans* function.

### Influence of cingulate sulcal morphology on MFC activation peaks

We hypothesized that the presence of a paracingulate sulcus in the medial frontal cortex would result in a shift of task activation peaks in the medial frontal cortex. To test this hypothesis, we performed a general linear model analysis to predict the MNI *z* -values of the medial frontal activation peaks with *cingulate sulcal morphology* and *task contrast* as predictors: 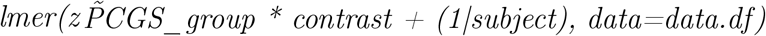 (Table 5)

**Table 5:** Model b: lmer(z hemisphere * contrast + (1|subject), data=data.df)

|  | Estimate | Std.Error | df | T | Pr(>\|z\|) |
| --- | --- | --- | --- | --- | --- |
| (Intercept) | 43.8774 | 0.6609 | 643.0000 | 66.389 | < 2e-16 *** |
| PCGS.L | 2.1176 | 1.1208 | 643.0000 | 1.889 | 0.05930 . |
| contrast.L | -4.2220 | 1.1557 | 643.0000 | -3.653 | 0.00028 *** |
| PCGS.L:contrast.L | 2.1719 | 1.9560 | 643.0000 | 1.110 | 0.26725 |

With this model, we observed a significant fixed main effect of task contrast (Table 5) : Going from basic movement control to movement monitoring to speech error monitoring, the average z-values of the activation peaks in the MFC decreased (i.e. shifted more ventrally; Figure 8). In line with our hypothesis, we observed a marginally significant effect (*p* = 0.0593) of cingulate sulcal morphology on the z-values of the medial frontal cortical activation peaks ((Table 5) : Going from subjects with no PCGS to those with an emerging PCGS and to those with a prominent PCGS, the mean z-values of the MFC activation peaks increase indicating that there was a dorsal shift in the MFC activation peaks with the presence of a PCGS (Figure 8.

**Figure 8:**
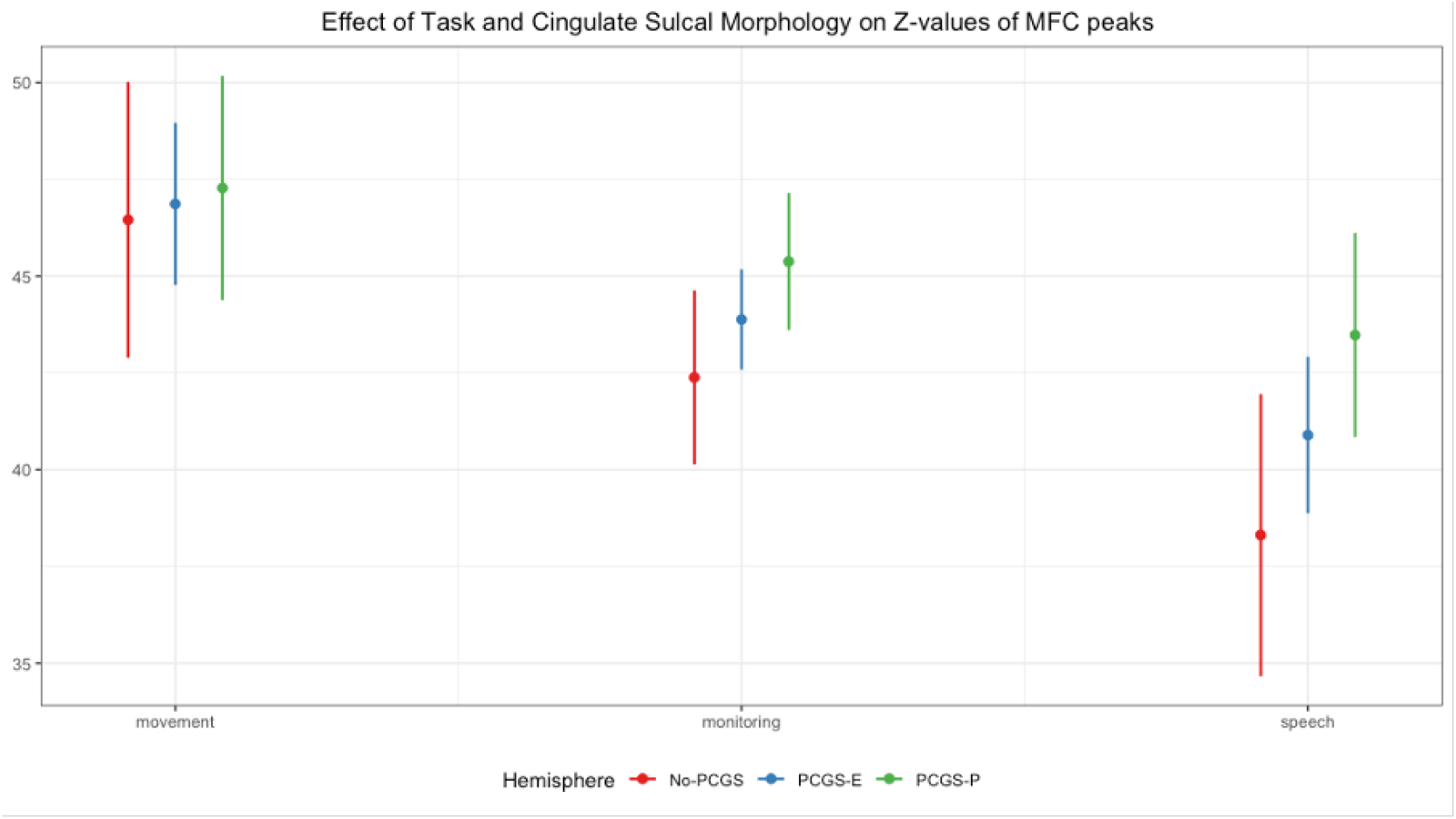
Predicted z-values of MFC activation peaks plotted by PCGS and task condition. Predicted peak probabilities were obtained via the fitting of model b : lmer(z PCGS * contrast + (1|subject), data=data.df), followed by the *ggmmeans* function.

Based on the Figure 8, the effect of cingulate sulcal morphology appeared more evident in the monitoring and speech task conditions. We performed a follow-up analysis where we fitted general linear models to predict z-values from the movement, monitoring and speech task conditions separately with cingulate sulcal morphology. These models revealed that the effect of cingulate sulcal morphology was significant only for the speech monitoring condition (*p* = 0.0377) but not for movement (*p* = 0.685) and monitoring conditions (*p* = 0.445).

## 8. Discussion

The main objective of the current study was to test the hypothesis that speech error monitoring is part of a more general action outcome evaluation computation performed in the medial frontal cortex that is distributed along a caudal-to-rostral, motor-to-cognitive control processing gradient. In a first study, we performed a fine-grained, individual-specific characterization of the medial frontal cortical activations related to speech error monitoring. We assessed the overlap of speech error monitoring activations with previously reported movement, feedback processing and cognitive control related activations in the medial frontal cortex and tested whether speech error monitoring activations are modulated by cingulate sulcal morphology (i.e. the presence or absence of a paracingulate sulcus) as previously observed for movement. We identified three activation clusters in the medial frontal cortex that are associated with speech error monitoring: one in the preSMA, and two peaks in the mid-cingulate cortex region (MCC1 and MCC2), which significantly overlapped with previously documented medial frontal activations for movement, feedback processing, and cognitive control. Additionally, the observed dorsal shift of the medial frontal activation peaks, in particular the posterior mid-cingulate cortex peak (i.e. MCC1) in the presence of a PCgS replicated findings from movement studies, suggesting a shared functional origin. In a second crucial study, we investigated, in individual subjects, the location of activation peaks in the medial frontal cortex that are associated with three task conditions that vary in the level of action control demands, namely basic tongue movements (motor control), tongue movements with monitoring (motor and cognitive control), and speech error monitoring (cognitive control). At the group level, we observed that all three task conditions elicited overlapping clusters of activation in the medial frontal cortex. However, by performing individual-level analysis of the medial frontal cortical activation peaks associated with the three task conditions, we were able to uncover fine-grained differences in medial frontal cortex activation patterns across the three task types: First, we found that with increasing cognitive control demands (i.e. going from the movement to the monitoring and to the speech error monitoring task) there is a higher probability of observing activation peaks in more rostral parts of the medial frontal cortex. Second, we observed that increasing cognitive control demands also resulted in a ventral shift in the medial frontal activation peaks. Another critical finding from our study was the demonstration that there was a significant effect of cingulate sulcal morphology, i.e. the presence or absence of a paracingulate sulcus, on the medial frontal activations associated with speech error monitoring. This effect, however, did not reach statistical significance for the movement and movement monitoring conditions. The last key finding was that across all three tasks, there was a higher prevalence of medial frontal activation peaks on the left versus right hemisphere. While leftward brain lateralisation was expected for speech error monitoring, the observation of leftward lateralisation for basic tongue movement control and the monitoring of tongue movements was highly intriguing.

### Speech error monitoring as part of a caudal-to-rostral gradient of action outcome evaluation in the medial frontal cortex

Our findings support the hypothesis that speech error monitoring is part of a broader action outcome evaluation computation. We observed that an overlapping set of regions in the medial frontal cortex were engaged across the basic and cognitive monitoring of tongue movements as well as speech error monitoring. However, individual-level analyses of medial frontal cortex activation peaks revealed a caudal-to-rostral shift of activations going from more basic to higher levels of behavioral monitoring. This rostral-caudal gradient for control demands aligns with Zarr and Brown (2016) proposal (2016), where prediction errors of increasing abstractness engage more rostral medial frontal cortex regions, such as the mid-cingulate area BA32 and the medial prefrontal areas BA8 and BA9. In our contrasts, prediction errors encoded in the caudal medial frontal cortex regions, such as the SMA and the preSMA, likely arise from discrepancies between sensory goals and states (e.g., Floegel et al. (2023)), while prediction errors encoded in the rostral medial frontal cortex result from mismatches between epistemic goals and states (e.g., levels of certainty, Pezzulo (2018); Pezzulo et al. (2022)). Alternatively, the rostral-caudal gradient might be interpreted in terms of accuracy, as proposed by Bonini et al. (2014). This would still be most parsimoniously conceived as a scalar variable of prediction error, though the relevant gradient dimension would be the magnitude of prediction error (low or high) rather than its type (sensory or epistemic). Future studies could parametrically and orthogonally vary motor and cognitve control as well as the absence or presence of overt errors to further support one of the two possibilities. Regardless of the exact variables driving the gradient, the prediction error framework would integrate actions of both motor and cognitive types, and thus both sensory and epistemic prediction errors. This would provide a unified explanation for previous apparently conflicting findings as both sensory feedback and conflict, re-characterized as sensory and epistemic sources of prediction error, could allow for evaluation of action outcomes through a single computation. In the case of speech error monitoring, both sensory and epistemic prediction error could be present, as speech production involves both sensory and epistemic goals. Concerning sensory feedback as a source of prediction error for speech errors, the influence of sulcal morphology on functional activations related to speech errors observed here resembles the pattern reported in prior research by Loh et al. (2020) for facial movement and feedback processing across various modalities. Assuming functional overlap, our findings suggest that sensory feedback from speech errors leads to mid-cingulate cortex activation due to a mismatch with desired sensory goals, signaling a need for adaptive control. However, the exact nature of this feedback provided through the speech errors remains an open question. The most apparent option is auditory feedback, but compelling evidence shows many speech errors are detected before articulation begins. For instance, when examining the latencies between the initiation of speech errors and their interruption, they exhibit a bimodal distribution: the interruption of an erroneous segment occurs either shortly after the onset of an error or approximately 500 milliseconds later (Nooteboom and Quené (2017)). Moreover, several studies using EEG have shown a modulation of the error-related negativity (ERN) for speech errors (e.g. Ganushchak and Schiller (2008); Riès et al. (2011); Baus et al. (2020), Dorokhova et al. (in press)). This ERP component has been source localized to the SMA/MCC and peaks around 100 ms after the onset of a response, rendering an explanation in terms of auditory feedback implausible. Thus, if the SMA/MCC activation from speech errors is feedback-related, proprioception is a more plausible source. Consistent with this interpretation, previous studies have shown the MFC is implicated in muscle-spindle feedback control (e.g., Goble et al. (2011)). Proprioceptive feedback control aligns with models integrating somatosensory targets to guide speech production (Guenther and Hickok (2016);Hickok (2012); Hickok (2014)) and recent findings showing speech errors are decoded above chance in a pre-response time window coinciding with the previously reported electromyographic speech-related activity (Dorokhova et al. (in press); Riès et al. (2012)). Concerning epistemic prediction error for speech production, one possibility would be to consider that response conflict would lead to changes in epistemic states deviating from the desired state, such as level of uncertainty, prior to error commission. This notion could be compatible with discussions in Botvinick et al. (2001)’s work proposing that conflict might constitute a component of a broader system primarily focused on reallocating attentional resources. In this scenario, conflict stemming from the competition between alternatives, feedback indicating error commission, and the experience of pain (e.g., Jones et al. (1991)), all of which have been shown to activate the MCC, collectively fall into the same category of cues signifying an insufficiency in the current allocation of attention resources to avert unfavorable outcomes, namely deviations from sensory or epistemic goals.

An intriguing possibility is that this caudal-rostral gradient in the medial frontal cortex indicate that this region encodes both control type, from motor to cognitive, along a parallel rostral-caudal gradient in the lateral frontal cortex, and representation type, from sensory to mental, along a parallel anterior-posterior gradient that is also observed in the parietal cortex, going from primary sensory areas to associative areas (e.g., Choi et al. (2018)). This idea aligns with the notion that the medial frontal cortex instructs frontal regions that control parietal representations. The prediction error signals that the medial frontal cortex detects could be generated either via comparisons between internal goals in frontal areas with parietal representations in initial learning stages and upon unexpected environmental changes, or via cerebellar internal models mimicking parietal representation dynamics (Ito (2008)). Consistent with this network view of adaptive control, previous studies examining speech errors and response conflict in language production have reported an involvement of the cerebellum as well as of frontal and parietal regions on top of the medial frontal recruitment already discussed (Runnqvist et al. (2016); Runnqvist et al. (2021); Todorović et al. (2023)). Outside of the domain of language, there is evidence that the cerebellum is organized along a gradient sensitive to executive control of different degrees of abstraction, similarly to frontal and parietal regions (D’Mello et al. (2020)). Moreover, while the cerebellar and cortical gradients were correlated, cortical gradients could not account for more than a small portion of the cerebellar gradients, suggesting that they have distinct functions as would be expected in the adaptive, internal modeling control network. Future research could use functional connectivity analyses to explore the distinct functions of nodes in this proposed adaptive control network.

### A leftward brain lateralisation for the control of basic tongue movements, tongue monitoring and speech error monitoring

Another interesting finding of our study was a significant prevalence of medial frontal activation peaks in the left hemisphere across the three task conditions. While it is expected for speech processing functions to be left-lateralised, the observation that the motor and cognitive control over basic tongue movements evoke more medial frontal cortex peaks in the left versus the right hemisphere suggested that non-linguistic and non-verbal tongue control is also specialised in the left hemisphere. Contradicting our findings, previous investigations about the neural correlates of tongue movements using functional MRI have largely revealed bilateral activations in the medial frontal cortex (Corfield et al. (1999); He et al. (2003); Martin et al. (2004); Amiez and Petrides (2014)). One possible explanation is that these studies (except for Amiez and Petrides (2014)) have performed group-level fMRI analyses and have relied on standard brain atlases to define the various medial frontal brain regions. This approach do not provide the fine-grained dissociations between the various medial frontal areas that is afforded by the individual-subject level analyses performed in the current study. While Amiez and Petrides (2014) have identified individual activation peaks that are associated with tongue movements in the medial frontal region, they did not analyse the hemispheric differences in these tongue movement related medial frontal activation peaks.

### A dorsal shift in medial frontal activations in relation to cingulate sulcal morphology

Previous research have revealed that the presence of a paracingulate sulcus in the medial frontal cortex results in a dorsal shift in the location of cytoarchitectonic area BA32 from the dorsal bank of the cingulate sulcus (if a paracingulate sulcus was not present) on to the paracingulate gyrus that is situated above the cingulate sulcus (Vogt et al. (1995)). Accompanying this dorsal shift in BA32, other studies have reported a corresponding dorsal shift in functional activations in the medial frontal cortex when a paracingulate sulcus was present (Loh et al. (2020); Loh et al. (2018); Amiez et al. (2006)). Adding to this body of work, we show that the location speech error monitoring peaks in individual brains also shift dorsally in the presence of a paracingulate sulcus. This finding underscores the importance of taking into account cingulate sulcal morphology when investigating the locations of speech processing in the medial frontal cortex.

### A ventral shift in medial frontal activations with increasing cognitive control demands

Turning to the ventral shift in the location of medial frontal activation peaks going from basic movement to movement monitoring and to speech error monitoring, this result might be partly explained by the fact that as the activations shift from more caudal (i.e. preSMA and SMA) to more rostral parts of the medial frontal cortex (i.e. mid-cingulate area 32 and medial prefrontal areas BA8, BA9), there is a decrease in z-values as the more rostral regions tend to occupy a more ventral position in the medial frontal cortex. Based on the cytoarchitectonic organisation of the cortical areas in the medial frontal region, Pandya et al. (2015) proposed that a ventral-to-dorsal gradient of architectonic differentiation exists in the medial frontal cortex starting from the indusium griseum, a primitive archicortical region that is located just above the corpus callosum; followed dorsally by the cingulate cortex (i.e. cortical areas BA32 and BA24), a proisocortical region which is a more differentiated cortex; and finally, the cortex on the superior frontal gyrus (i.e. SMA and preSMA (BA6), and medial prefrontal areas BA8 and BA9) which is the phylogenetically newer six-layered isocortex. Barbas (2015) further suggested that less differentiated parts of the frontal cortex are more strongly innervated by subcortical structures that are involved in emotional and motivational processing, such as the thalamus, amygdala and the hippocampus, and they send feedback-type connections to regulate the activity in more differentiated frontal areas for flexible cognition and behaviors. Based on this anatomical account, the ventral shift in medial frontal activations with increasing cognitive control demands could reflect the increased recruitment of the more ventral limbic-related frontal brain regions for more complex behavioral control.

Finally, our findings offer implications beyond fundamental research. For instance, in neurosurgery, precise brain region localization is imperative. Identifying potential anatomical landmarks within individual brains, associated with functions from movement control to speech monitoring, could help surgeons enhance procedural precision and minimize collateral damage, improving patient outcomes. Beyond clinical applications, these findings provide researchers with invaluable references for investigating the neural substrates of speech production and error monitoring, promising fundamental insights into the mechanisms governing language production. Furthermore, our findings add to a growing body of research showing that sulcal morphology influences the functional organization of the human and non-human primate brain (e.g., Hopkins et al. (2017); Lopez-Persem et al. (2019); Leroy et al. (2015); Bodin et al. (2018); Eichert et al. (2021)). In conclusion, our study robustly supports the hypothesis that speech error monitoring is part of a broader action outcome evaluation mechanism, distributed along a medial frontal cortex (MFC) rostral-caudal processing gradient. We identified activation clusters ranging from the SMA to BA9m, which overlapped significantly with areas involved in movement, feedback processing, and cognitive control, further modulated by sulcal morphology. The linear increase in activation peaks with escalating cognitive control demands in the rostral direction underscores the MFC’s role in managing prediction errors of varying abstractness. Additionally, the unexpected dorsal-ventral gradient and left hemisphere prevalence of peaks suggest complex spatial organization of control processes, potentially indicating the MFC’s encoding of both control type and representation type. These insights not only advance our understanding of the neural substrates underlying speech production and error monitoring but also hold significant implications for clinical practices, such as neurosurgery, and broader cognitive neuroscience research. Future investigations should further dissect the distinct roles of various brain regions for motor and cognitive control, as well as in relation to the sources driving prediction errors.

## 9. Funding and acknowledgements

This work, carried out within the Institute of Convergence ILCB (ANR-16-CONV-0002), has benefited from support from the French government (France 2030), managed by the French National Agency for Research (ANR) and the Excellence Initiative of Aix-Marseille University (A*MIDEX). E.R. has benefited from support from the French government, managed by the French National Agency for Research (ANR) through a research grant (ANR-18-CE28-0013). The work was performed in the Center IRM-INT (UMR 7289, AMU-CNRS), platform member of France Life Imaging network (grant ANR-11-INBS-0006). Centre de Calcul Intensif d’Aix-Marseille is acknowledged for granting access to its high performance computing resources. The authors are grateful to Linda Dahmani for annotating part of the functional data.

## Appendix A. Tables

**Table A.1:** Individual activation peaks in the PreSMA region. The x, y, z, coordinates are in MNI stereotaxic space. Individual peaks are identified at a significance threshold of p<0.05, corrected for multiple comparisons within the medial frontal cortex.

| Subject | PCGS Group | Left |  |  |  | PCGS Group | Right |  |  |  |
| --- | --- | --- | --- | --- | --- | --- | --- | --- | --- | --- |
|  |  | *x* | *y* | *z* | *t* |  | *x* | *y* | *z* | *t* |
| 1 | Prominent | - | - | - | - | Absent | - | - | - | - |
| 2 | Prominent | -6 | 14 | 59 | 4.04 | Absent | 5 | 15 | 58 | 5.22 |
| 3 | Emerging | -4 | 9 | 57 | 3.14 | Absent | - | - | - | - |
| 4 | Emerging | -5 | 4 | 66 | 2.6 | Prominent | - | - | - | - |
| 5 | Prominent | -1 | 10 | 55 | 3.88 | Emerging | 3 | 10 | 54 | 3.6 |
| 6 | Prominent | - | - | - | - | Emerging | - | - | - | - |
| 7 | Emerging | - | - | - | - | Absent | - | - | - | - |
| 8 | Absent | - | - | - | - | Absent | - | - | - | - |
| 9 | Emerging | -4 | 13 | 63 | 3.23 | Absent | - | - | - | - |
| 10 | Emerging | - | - | - | - | Prominent | - | - | - | - |
| 11 | Emerging | -8 | 16 | 58 | 4.18 | Absent | 3 | 11 | 55 | 3.84 |
| 13 | Emerging | -2 | 14 | 65 | 6.67 | Emerging | - | - | - | - |
| 14 | Emerging | - | - | - | - | Prominent | - | - | - | - |
| 16 | Absent | -5 | 10 | 52 | 8.28 | Absent | 3 | 8 | 54 | 6.94 |
| 17 | Absent | -11 | 7 | 60 | 2.42 | Absent | - | - | - | - |
| 19 | Absent | - | - | - | - | Absent | - | - | - | - |
| 20 | Emerging | - | - | - | - | Absent | - | - | - | - |
| 21 | Emerging | -2 | 7 | 57 | 6.97 | Emerging | 5 | 14 | 65 | 6.4 |
| 22 | Prominent | -8 | 10 | 58 | 2.2 | Prominent | - | - | - | - |
| 23 | Prominent | -2 | 16 | 53 | 2.22 | Prominent | 7 | 17 | 60 | 3.29 |
| 25 | Prominent | -7 | 8 | 53 | 2.21 | Prominent | - | - | - | - |
| 26 | Emerging | - | - | - | - | Absent | - | - | - | - |
| 27 | Prominent | -3 | 8 | 67 | 3.69 | Emerging | 3 | 8 | 65 | 3.55 |
| 28 | Absent | - | - | - | - | Absent | - | - | - | - |

**Table A.2:**
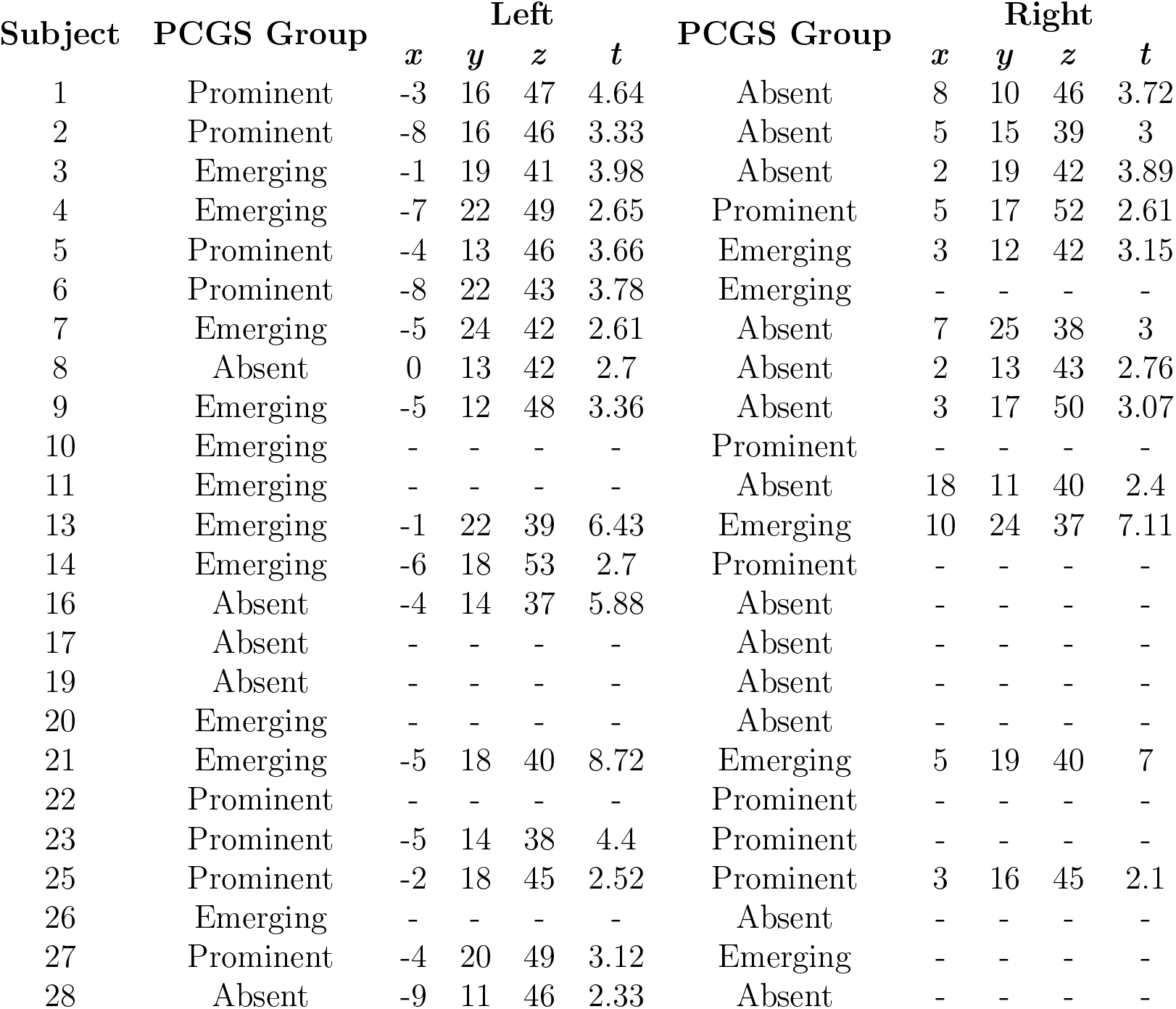
Individual activation peaks in the posterior MCC (MCC1). The x, y, z, coordinates are in MNI stereotaxic space. Individual peaks are identified at a significance threshold of p<0.05, corrected for multiple comparisons within the medial frontal cortex.

| Subject | PCGS Group | Left |  |  |  | PCGS Group | Right |  |  |  |
| --- | --- | --- | --- | --- | --- | --- | --- | --- | --- | --- |
|  |  | *x* | *y* | *z* | *t* |  | *x* | *y* | *z* | *t* |
| 1 | Prominent | -3 | 16 | 47 | 4.64 | Absent | 8 | 10 | 46 | 3.72 |
| 2 | Prominent | -8 | 16 | 46 | 3.33 | Absent | 5 | 15 | 39 | 3 |
| 3 | Emerging | -1 | 19 | 41 | 3.98 | Absent | 2 | 19 | 42 | 3.89 |
| 4 | Emerging | -7 | 22 | 49 | 2.65 | Prominent | 5 | 17 | 52 | 2.61 |
| 5 | Prominent | -4 | 13 | 46 | 3.66 | Emerging | 3 | 12 | 42 | 3.15 |
| 6 | Prominent | -8 | 22 | 43 | 3.78 | Emerging | - | - | - | - |
| 7 | Emerging | -5 | 24 | 42 | 2.61 | Absent | 7 | 25 | 38 | 3 |
| 8 | Absent | 0 | 13 | 42 | 2.7 | Absent | 2 | 13 | 43 | 2.76 |
| 9 | Emerging | -5 | 12 | 48 | 3.36 | Absent | 3 | 17 | 50 | 3.07 |
| 10 | Emerging | - | - | - | - | Prominent | - | - | - | - |
| 11 | Emerging | - | - | - | - | Absent | 18 | 11 | 40 | 2.4 |
| 13 | Emerging | -1 | 22 | 39 | 6.43 | Emerging | 10 | 24 | 37 | 7.11 |
| 14 | Emerging | -6 | 18 | 53 | 2.7 | Prominent | - | - | - | - |
| 16 | Absent | -4 | 14 | 37 | 5.88 | Absent | - | - | - | - |
| 17 | Absent | - | - | - | - | Absent | - | - | - | - |
| 19 | Absent | - | - | - | - | Absent | - | - | - | - |
| 20 | Emerging | - | - | - | - | Absent | - | - | - | - |
| 21 | Emerging | -5 | 18 | 40 | 8.72 | Emerging | 5 | 19 | 40 | 7 |
| 22 | Prominent | - | - | - | - | Prominent | - | - | - | - |
| 23 | Prominent | -5 | 14 | 38 | 4.4 | Prominent | - | - | - | - |
| 25 | Prominent | -2 | 18 | 45 | 2.52 | Prominent | 3 | 16 | 45 | 2.1 |
| 26 | Emerging | - | - | - | - | Absent | - | - | - | - |
| 27 | Prominent | -4 | 20 | 49 | 3.12 | Emerging | - | - | - | - |
| 28 | Absent | -9 | 11 | 46 | 2.33 | Absent | - | - | - | - |

**Table A.3:** Individual activation peaks in the anterior MCC (MCC2). The x, y, z, coordinates are in MNI stereotaxic space. Individual peaks are identified at a significance threshold of p<0.05, corrected for multiple comparisons within the medial frontal cortex.

| Subject | PCGS Group | Left |  |  |  | PCGS Group | Right |  |  |  |
| --- | --- | --- | --- | --- | --- | --- | --- | --- | --- | --- |
|  |  | *x* | *y* | *z* | *t* |  | *x* | *y* | *z* | *t* |
| 1 | Prominent | -7 | 24 | 28 | 2.8 | Absent | 7 | 25 | 28 | 3.23 |
| 2 | Prominent | -3 | 29 | 33 | 4.33 | Absent | 9 | 33 | 26 | 6.47 |
| 3 | Emerging | -1 | 28 | 15 | 3.17 | Absent | 8 | 29 | 30 | 3.7 |
| 4 | Emerging | - | - | - | - | Prominent | - | - | - | - |
| 5 | Prominent | -10 | 30 | 17 | 2.87 | Emerging | 12 | 25 | 39 | 2.7 |
| 6 | Prominent | -1 | 29 | 32 | 4.93 | Emerging | 3 | 29 | 32 | 4.86 |
| 7 | Emerging | -9 | 24 | 34 | 2.42 | Absent | 8 | 29 | 31 | 3.38 |
| 8 | Absent | -2 | 18 | 33 | 2.75 | Absent | 3 | 24 | 26 | 2.87 |
| 9 | Emerging | -6 | 29 | 40 | 2.38 | Absent | 9 | 21 | 35 | 2.3 |
| 10 | Emerging | -1 | 30 | 43 | 2.37 | Prominent | 4 | 31 | 40 | 2.54 |
| 11 | Emerging | -5 | 28 | 28 | 3.91 | Absent | 12 | 26 | 29 | 4.31 |
| 13 | Emerging | -9 | 34 | 20 | 6.61 | Emerging | 3 | 24 | 21 | 3.62 |
| 14 | Emerging | -6 | 25 | 40 | 2.29 | Prominent | - | - | - | - |
| 16 | Absent | -5 | 23 | 28 | 4.62 | Absent | 8 | 22 | 39 | 6.71 |
| 17 | Absent | - | - | - | - | Absent | - | - | - | - |
| 19 | Absent | - | - | - | - | Absent | - | - | - | - |
| 20 | Emerging | - | - | - | - | Absent | - | - | - | - |
| 21 | Emerging | - | - | - | - | Emerging | 10 | 26 | 25 | 3.84 |
| 22 | Prominent | - | - | - | - | Prominent | - | - | - | - |
| 23 | Prominent | -1 | 27 | 32 | 5.36 | Prominent | 10 | 28 | 33 | 5.99 |
| 25 | Prominent | - | - | - | - | Prominent | 8 | 27 | 27 | 2.03 |
| 26 | Emerging | - | - | - | - | Absent | - | - | - | - |
| 27 | Prominent | -6 | 29 | 26 | 4.09 | Emerging | 7 | 23 | 32 | 3.6 |
| 28 | Absent | - | - | - | - | Absent | 8 | 29 | 27 | 2.91 |

**Table A.4:** Coordinates, *t* values and cluster sizes of peaks of activation in left hemisphere across three groups outside the area of main interest (ACC). Significant *t* values for exploratory search are displayed in bold text.

|  | cluster size | t | MNI coordinates |  |  |
| --- | --- | --- | --- | --- | --- |
|  |  |  | x | y | z |
| Prominent PCgS | 242 | **13.43** | **-32** | **26** | **-10** |
|  |  | **8.50** | **-52** | **9** | **-2** |
|  |  | 7.30 | -55 | 12 | 8 |
|  | 13 | **9.65** | **-5** | **-4** | **3** |
|  | 10 | **7.59** | **-42** | **-4** | **53** |
|  | 2 | 5.92 | -45 | 14 | 20 |
| Emerging PCgS | 13 | **6.95** | **-40** | **9** | **26** |
|  | 15 | 6.58 | -35 | 29 | -2 |
|  | 12 | 6.47 | -10 | -6 | 6 |
|  | 6 | 6.01 | -42 | -1 | 48 |
|  | 1 | 5.86 | -15 | -4 | 10 |
|  | 1 | 5.62 | -52 | 6 | 38 |
|  | 1 | 5.44 | -42 | -38 | 46 |
|  | 2 | 5.43 | -42 | 24 | -10 |
| Absent PCgS | 1 | **6.74** | **-2** | **-14** | **-12** |
|  | 32 | 6.50 | -38 | 26 | 3 |
|  |  | 6.41 | -32 | 22 | 10 |

## Appendix B. Figures

**Figure B.1:**
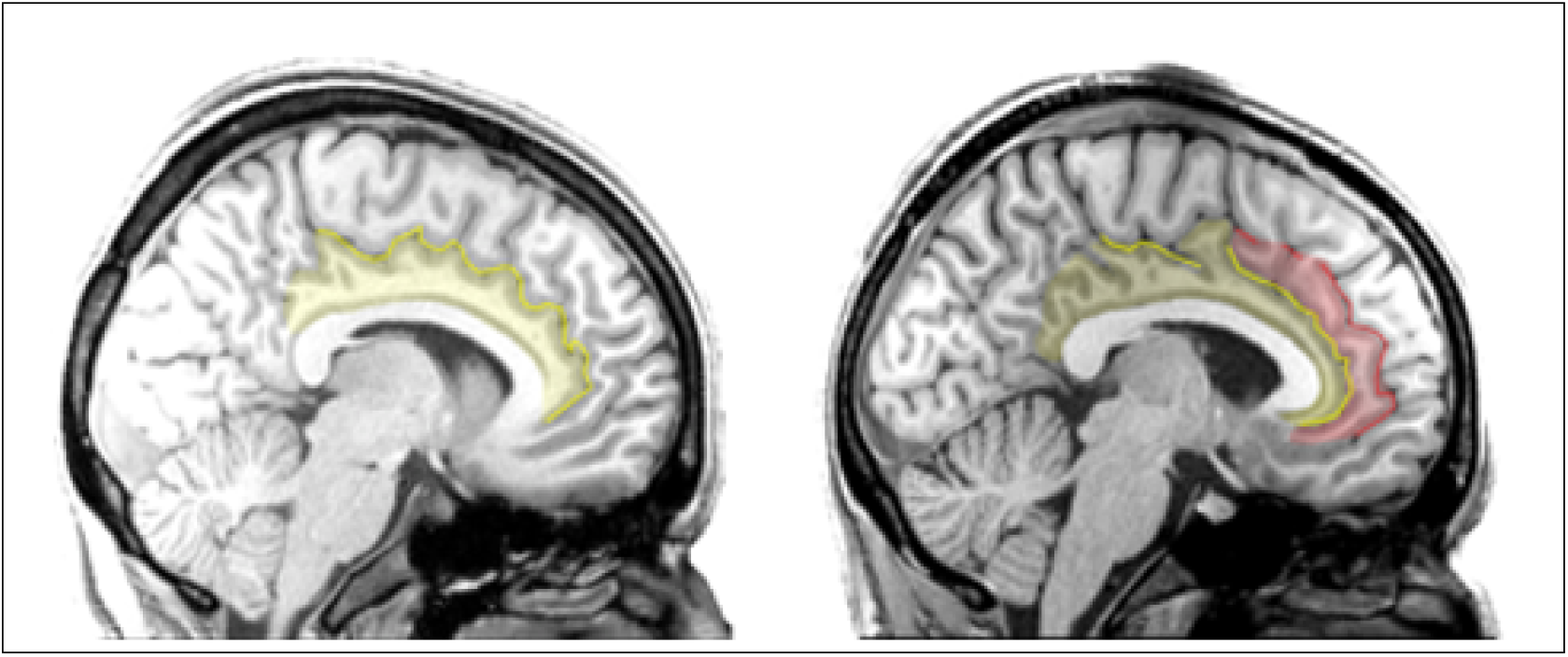
Examples of a hemisphere with only one CgS and a cingulate gyrus (left) and a hemisphere with a CgS and a prominent PCgS and paracingulate gyrus (right). Sulci are displayed with lines of different colors: yellow for the CgS, red for the PCgS; the corresponding color shades overlay the corresponding gyri.

**Figure B.2:**
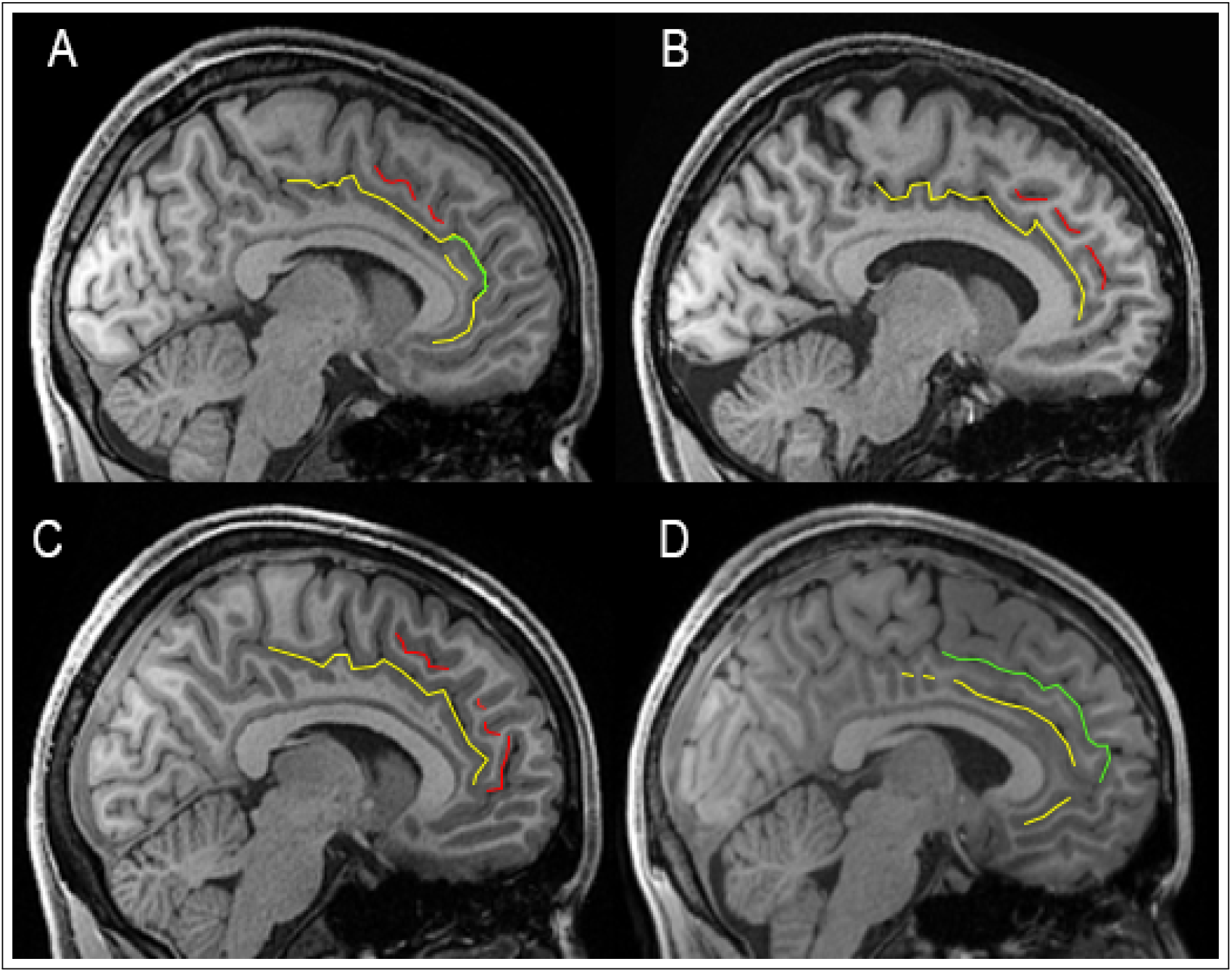
Examples of several emerging cases. The CgS is always displayed in yellow color, the possible PCgS is displayed in red and sulci that could be both CgS and PCgS are in green. The two examples of the top panels are left hemispheres and the two examples of the bottom panels are right hemispheres. Reasons for the classification as “Emerging”: A -short and discontinued, B-discontinued, C - discontinued, D - not enough lateral depth.

**Figure B.3:**
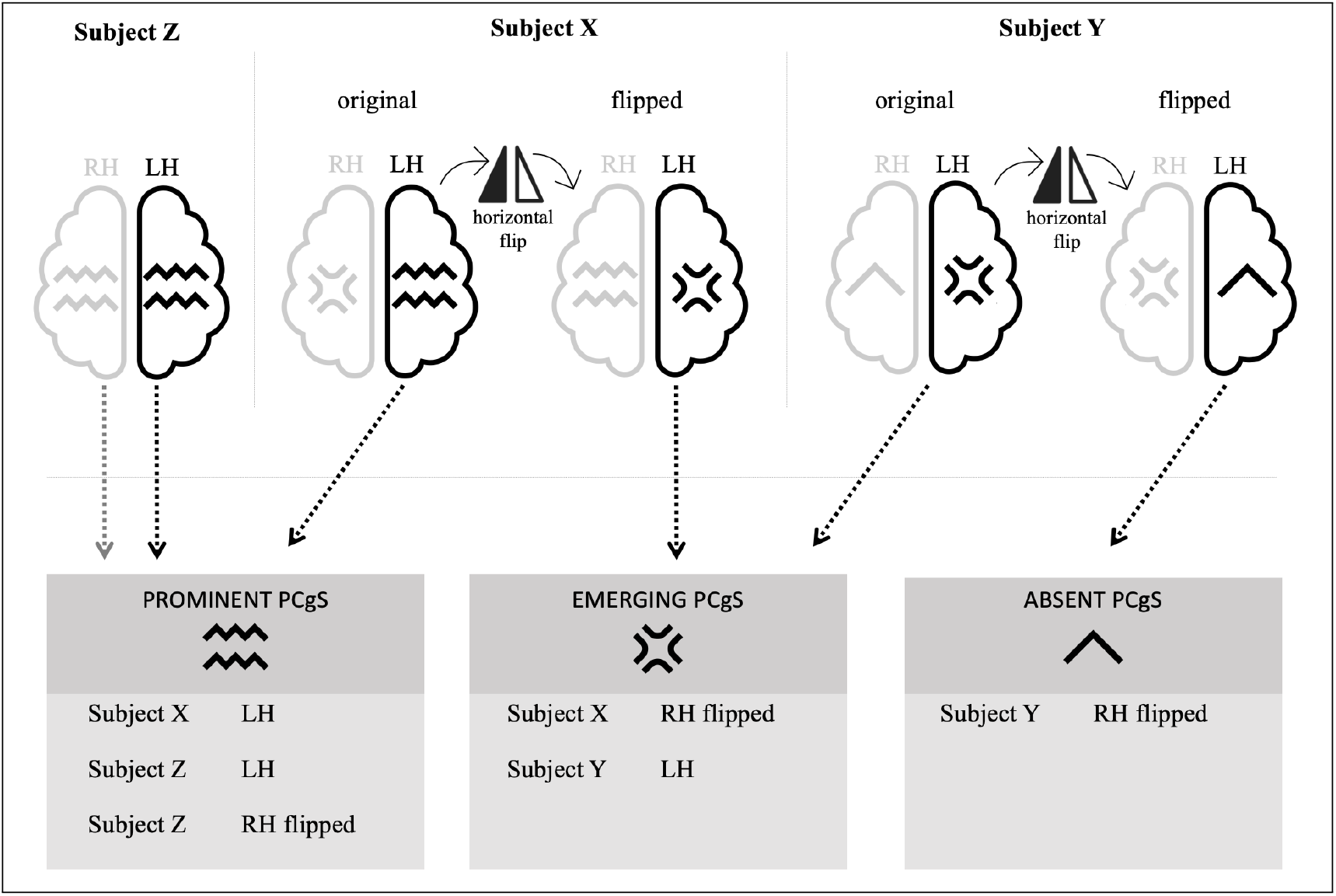
A schematic representation of the process of flipping and dispatch into the groups. RH - right hemisphere, LH - left hemisphere. The flipping procedure was done on T1-weighted scans as well as on functional contrast maps.

**Figure B.4:**
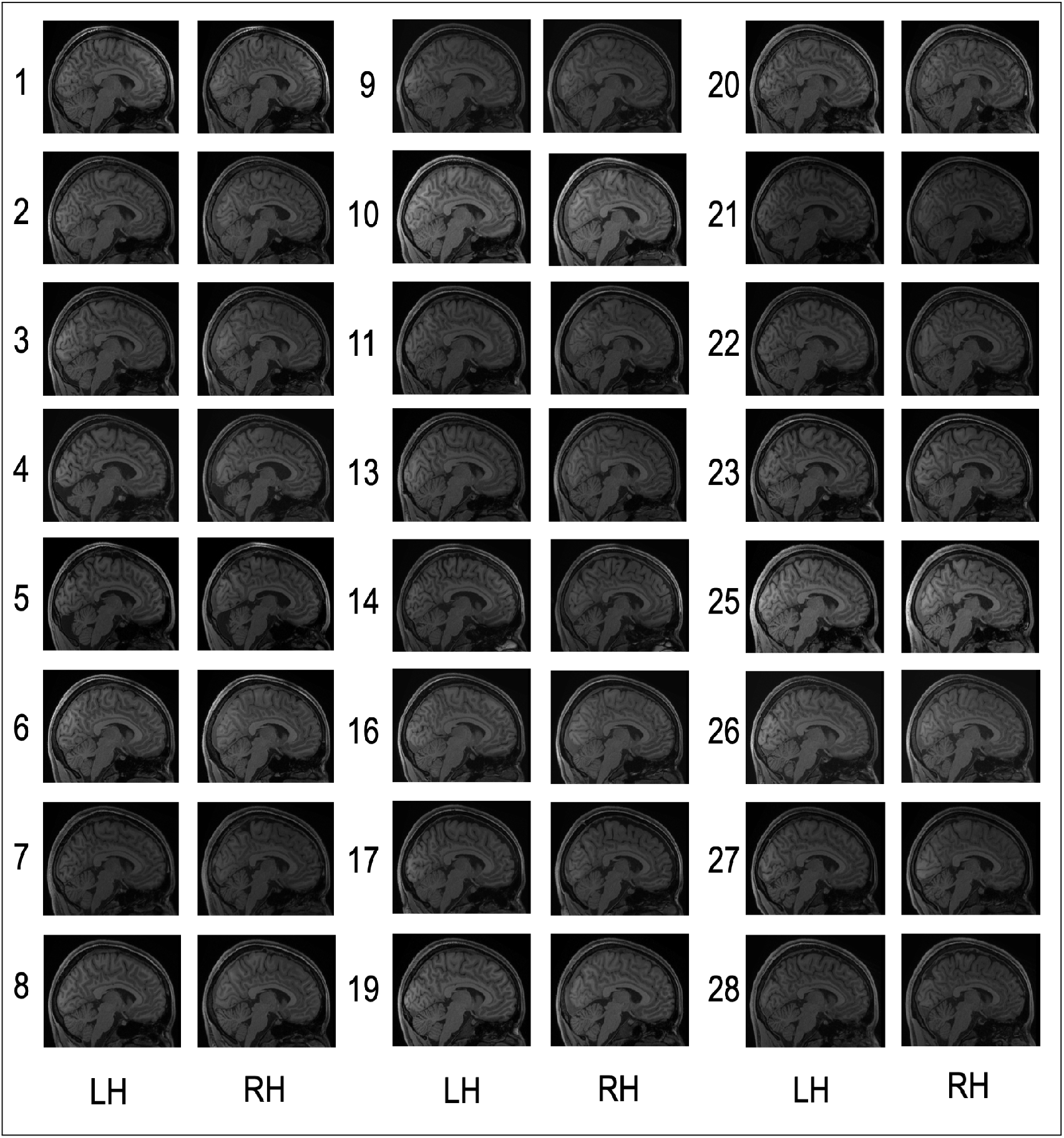
Sagittal slices of right (RH) and left (LH) hemispheres of all analysed subjects at *x* = 5 and *x* = −5 respectively.

## Notes

### Competing Interest Statement

The authors have declared no competing interest.

